# Counterfactual effect of Colombian protected areas on multispecies functional connectivity

**DOI:** 10.64898/2026.10.13.757925

**Authors:** Néstor Raúl Espejo-Delgado, Adriana Rojas-Suárez, David Santiago Afanador-Rivera, Nelly Rodríguez-Eraso

**Affiliations:** Facultad del Medio Ambiente y Recursos Naturales, Universidad Distrital Francisco José de Caldas, Bogotá, Colombia; Parques Nacionales Naturales de Colombia, Subdirección de Gestión y Manejo de Áreas Protegidas, Bogotá, Colombia; Fundación Gaia Amazonas, Bogotá, Colombia; Departamento de Biología, Universidad Nacional de Colombia, Bogotá, Colombia

**Keywords:** avoided loss, circuit theory, emergent functional groups, human modification, Kunming–Montreal Global Biodiversity Framework, Neotropics, propensity score matching

## Abstract

Protected areas (PAs) are central to Target 3 of the Kunming–Montreal Global Biodiversity Framework, which calls for ecologically well-connected systems, yet progress is tracked with indicators of connectivity state rather than of what protection achieved. We evaluated whether PAs in Colombia maintained multispecies functional connectivity relative to unprotected counterfactuals from 2000 to 2022 and whether the magnitude of their effects reflected the pressure they faced. We grouped 181 bird and mammal species into 12 emergent functional groups (EFGs) based on movement traits and exposure to human modification, modeled their connectivity with omnidirectional circuit theory, and matched protected and unprotected pixels with propensity scores within 6 regional subsystems. For 135 PAs, we estimated effects on standardized connectivity change (ΔZ) and on avoided critical loss (connected pixels that became impeded) and classified PAs by avoided loss relative to control loss. Connectivity changed more favorably in PAs than in controls (median ΔZ effect 0.87, 95% CI 0.36–1.60); 72.6% of PAs had positive effects, which were significant in 4 of 6 regions. Protected areas avoided 1.51 percentage points (pp) of critical loss nationally, and avoided loss was positive in all 12 EFGs and significant in 8. However, the presence of an effect overstated its contribution: only 55% of PAs avoided more than 1 pp, and 30% lost more than their controls. Effectiveness increased with pressure, from 39% of PAs under low pressure to 75% under high pressure, where sustainable-use areas accounted for 29 of the 33 effective PAs. Small PAs contained most of the loss expected without protection (median 70%), and regional and national PAs performed alike under similar pressure. Measuring avoided loss relative to pressure shows where protection secures connectivity and where governance must improve, and offers a replicable indicator for the well-connected component of Target 3.

## 1. Introduction

Protected areas (PAs) represent the primary strategy for safeguarding biodiversity in the context of global change (Gurney et al., 2023; Rodrigues & Cazalis, 2020). However, considerable debate persists regarding their effectiveness in delivering conservation outcomes (Geldmann et al., 2013; Jones et al., 2018). The conservation value of PAs depends not only on their internal biodiversity but also on their degree of ecological connectivity with surrounding landscapes (Brennan et al., 2022). Ecological connectivity, defined as the unimpeded movement of species and the flow of natural processes that sustain life on Earth (CMS, 2024), is essential for maintaining gene flow, preventing population isolation, and enabling species to respond to environmental change (Brodie et al., 2025; Liczner et al., 2024). For this reason, PAs are most effective when functioning as core areas of broader ecological networks, interconnected by corridors and permeable landscapes that facilitate species movement (Hilty et al., 2020).

Ecological connectivity encompasses both a structural dimension, determined by the spatial configuration of habitats (Baudry & Merriam, 1988), and a functional dimension, defined by organism-specific movement through the landscape (Taylor et al., 1993). As surrounding land is increasingly transformed, functional connectivity may decline even if habitat within PAs remains intact (Brennan et al., 2022; Sacre et al., 2025). Because functional connectivity is species-specific, single-species models cannot represent how the many species that share a landscape move through it (Koen et al., 2014). Multispecies approaches address this by identifying networks that sustain movement for many organisms at once (Prima et al., 2024; Wood et al., 2022), often by grouping species into emergent functional groups (EFGs) based on habitat requirements and life-history traits (Lechner et al., 2017; Vos et al., 2001). EFGs reduce analytical dimensionality while retaining ecological realism (Brennan et al., 2022; Liczner et al., 2024). Birds and terrestrial mammals are the most common models, given the information available on their dispersal and responses to landscape transformation (C. Correa-Ayram et al., 2016; Dickson et al., 2019).

Connectivity has also become a global policy commitment. Target 3 of the Kunming–Montreal Global Biodiversity Framework (KM-GBF) commits countries to protect 30% of land and sea by 2030 in ecologically well-connected systems (CBD, 2022). Yet progress on this component is tracked with state-level indicators. Structural indices such as ProtConn estimate that less than 10% of the world’s land is both protected and connected (Saura et al., 2018, 2019), and functional models for terrestrial mammals show that many PAs are poorly connected (Brennan et al., 2022). These indicators describe how connected a network is at a given time, but not whether protection made a difference.

Determining the true effect of protection is a counterfactual question (Ferraro, 2009). A PA experiencing biodiversity decline may still be considered effective if it reduces pressures compared with a scenario without protection. Conversely, a PA that would not have lost biodiversity even in the absence of pressures cannot be deemed effective (Ferraro, 2009; Joppa & Pfaff, 2011). Since PAs are frequently situated far from roads and cities, often on steep or unproductive land, comparisons with adjacent areas tend to overestimate their effectiveness (Joppa & Pfaff, 2009).

Statistical matching addresses this bias by comparing protected sites with unprotected sites possessing similar characteristics (Andam et al., 2008; Schleicher et al., 2020). Matching-based evaluations have shown that PAs and other area-based interventions reduce deforestation and habitat loss and fragmentation, but with effects that vary widely among regions, management types, and levels of threat (Lessmann et al., 2024; Li et al., 2024; Liévano-Latorre et al., 2021; Negret et al., 2020), and that effects on biodiversity itself are mixed (Cazalis et al., 2020; Santangeli et al., 2023).

However, to date, statistical matching has not been applied to functional connectivity. Furthermore, the magnitude of an intervention’s effect depends on the pressure an area faces, and for functional connectivity it should also depend on how strictly the area is protected, on the regional system that contains it, and on how the species considered move. Interventions prevent greater absolute losses where threats are high, while relative reductions may be larger where threats are low (Joppa & Pfaff, 2011; Nelson & Chomitz, 2011). Therefore, the magnitude of an effect must be interpreted in relation to the specific pressures each PA encounters (Pressey et al., 2015; Pressey et al., 2021).

Colombia captures many of the conditions that make this question relevant worldwide. The country’s protected area system (SINAP) encompasses the full management spectrum, ranging from large, nationally managed strict-protection parks to small, regionally managed sustainable-use areas (DNP, 2021). Colombian regions include landscapes that have been transformed since colonial times as well as extensive natural ecosystems now facing new deforestation frontiers (Clerici et al., 2020; C. Correa-Ayram et al., 2017; Murillo-Sandoval et al., 2022). These conditions are shared by many tropical countries expanding their PA systems toward 30% coverage (Fajardo et al., 2026).

Colombian PAs reduce deforestation, but their effect varies with regional context (Bonilla-Mejía & Higuera-Mendieta, 2019; Forero-Medina et al., 2026; Liévano-Latorre et al., 2021; Negret et al., 2020), providing an opportunity to test whether ecological connectivity outcomes exhibit similar regional variation. The national PA policy aims to integrate the PA system into connectivity networks consistent with the KM-GBF (DNP, 2021; MADS, 2024). Connectivity studies in Colombia have mapped corridors and prioritized areas for conservation and restoration (Linero-Triana et al., 2023; Mosquera-Guerra et al., 2025; Pineda-Zapata et al., 2024; Zárrate-Charry et al., 2018), and recent evidence shows that connectivity is declining in and around PAs (Murillo-Sandoval et al., 2022; Rubiano et al., 2026). Whether protection slowed that decline, however, remains untested.

We evaluated whether PAs maintained multispecies functional connectivity compared with unprotected counterfactuals from 2000 to 2022 and whether the effect’s magnitude reflected the pressure each PA faced. We grouped terrestrial birds and mammals into emergent functional groups (EFGs), modeled their connectivity with omnidirectional circuit theory, and estimated effects at PA and system levels through statistical matching. We hypothesized that PAs would show more favorable connectivity trajectories than their counterfactuals, consistent with evidence on avoided deforestation (hypothesis 1). We also expected effects to vary with functional group, region, management category, and size (hypothesis 2). Finally, we predicted that considering the pressure in matched controls would reduce the share of effective PAs and narrow the apparent advantage of sustainable-use over strict PAs (hypothesis 3).

## 2. Methods

### 2.1. Study area

Colombia encompasses approximately 1.1 million km² of continental territory in northwestern South America. Its diverse topography, soils, and climate support exceptional species and ecosystem diversity (Rangel-Ch, 2015). The National System of Protected Areas (SINAP) serves as the primary territorial conservation strategy, covering about 20 million continental hectares, equivalent to 17% of the national territory (RUNAP, 2026). The system includes a management gradient ranging from large, nationally managed strict-protection areas such as National and Regional Parks (IUCN categories Ia, Ib, and II) to smaller, regionally managed sustainable-use areas such as Integrated Management Districts and Protective Forest Reserves (IUCN category V) (Bonilla-Mejía & Higuera-Mendieta, 2019; DNP, 2021). The system is organized into 6 regional subsystems (SIRAPs), defined by biogeographic and administrative criteria: Amazon, Northeastern Andes, Western Andes, Caribbean, Orinoco, and Pacific (Fig. 1).

**Figure 1.**
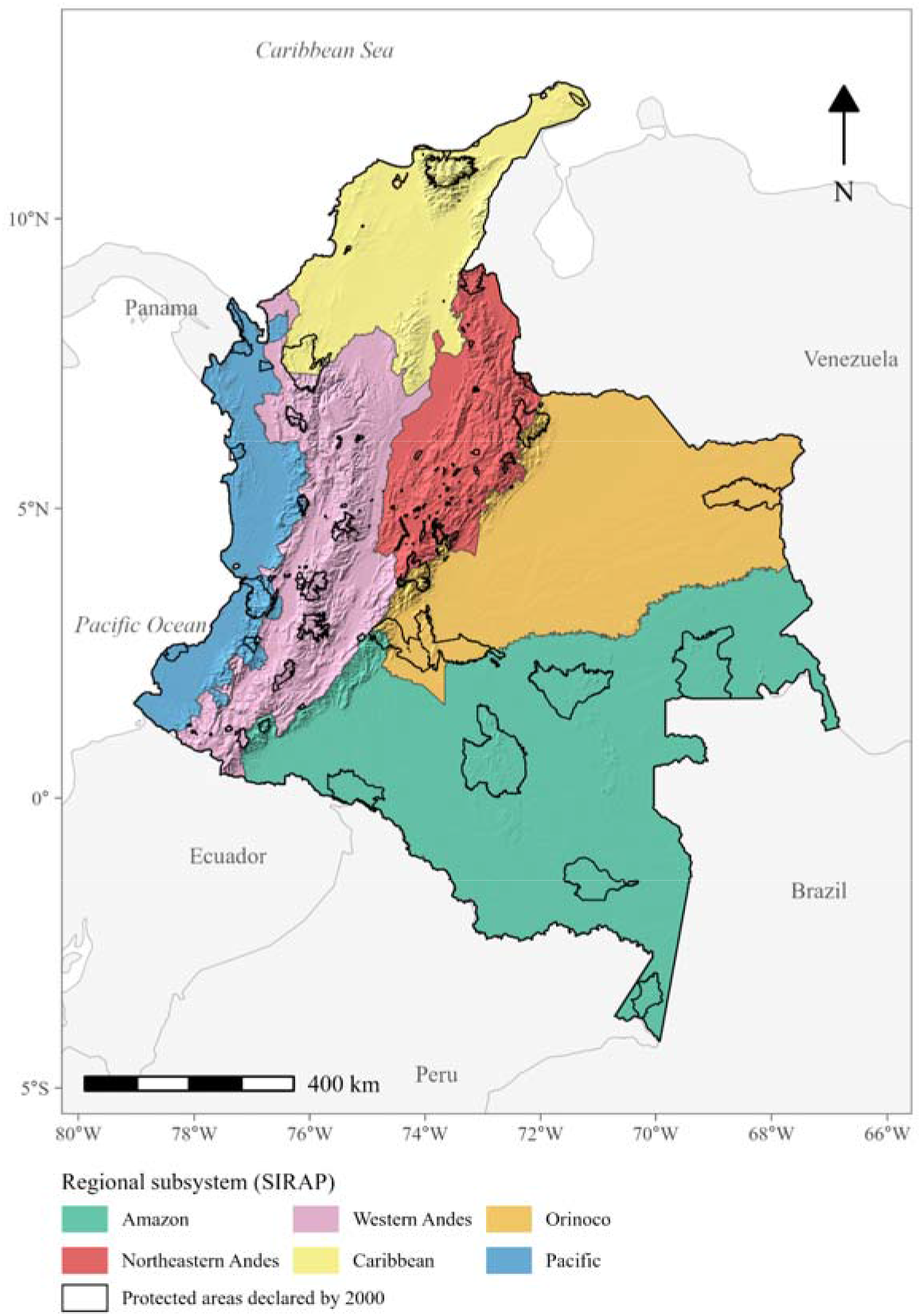
Study area. Regional subsystems of protected areas (SIRAP) of Colombia’s National System of Protected Areas (SINAP): Amazon, Northeastern Andes, Western Andes, Caribbean, Orinoco, and Pacific. Black outlines show the 135 protected areas declared by 2000 and included in the analysis.

To evaluate PA effects over the full study period, we considered only PAs declared by 2000 (Fig. 1). In that year, the system comprised 144 PAs covering 100,842 km², or 8.8% of the continental territory, compared with 17% today. Pixels in PAs declared after 2000 were used neither as treatment nor as control. We obtained boundaries, declaration dates, management categories, and jurisdiction from the official national registry of protected areas (RUNAP, 2026), which documents the legal act and date of each declaration.

### 2.2. Study design

We organized the study into 3 phases: definition of EFGs of birds and mammals, omnidirectional connectivity modeling for each EFG in 2000 and 2022 at 300-m resolution, and counterfactual evaluation of PA effects with 2 indicators, standardized connectivity change (ΔZ) and critical loss transition (CLT) (Fig. 2). Rather than relying on a few focal or generic species, we grouped species with similar traits, which reduces analytical complexity while retaining differences in how species move (Prima et al., 2024; Wood et al., 2022).

**Figure 2.**
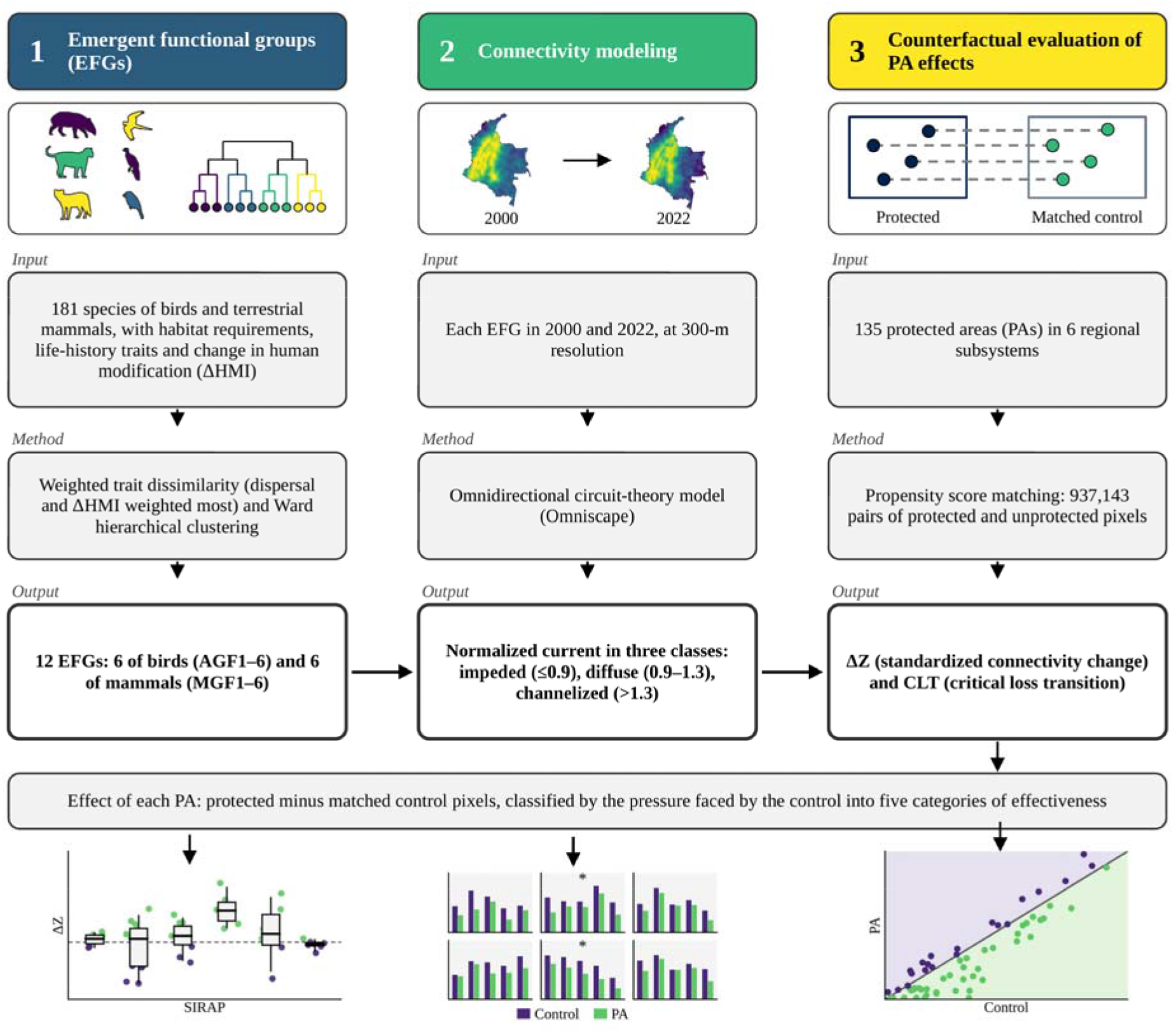
Conceptual framework of the study. (1) Species of birds and terrestrial mammals (181) were grouped by traits and change in human modification (ΔHMI) into 12 emergent functional groups (EFGs: AGF1–6 for birds, MGF1–6 for mammals). (2) Omnidirectional functional connectivity was modeled for each EFG in 2000 and 2022 at 300-m resolution with circuit theory (Omniscape), and normalized current was classified as impeded (≤0.9), diffuse (0.9–1.3), or channelized (>1.3). (3) The effect of each protected area (PA) was estimated as the difference between protected pixels and matched unprotected control pixels (propensity score matching; 937,143 pairs) with two indicators: the standardized connectivity change (ΔZ) and the critical loss transition (CLT). Effects were classified into five categories of effectiveness according to the pressure faced by the control. Illustrations are schematic and do not show data.

### 2.3. Delimitating emergent functional groups (EFGs)

We compiled an initial list of 348 terrestrial bird and mammal species from a systematic review of multispecies connectivity studies in the Neotropics. These taxa have the most complete information on dispersal and responses to landscape change (Correa-Ayram et al., 2016; Dickson et al., 2019). We excluded bats, for which few Neotropical dispersal estimates were available; aquatic and semiaquatic mammals; and aerial birds (*e.g.*, swifts, swallows, hummingbirds, and vultures), whose movements are poorly represented by terrestrial resistance surfaces (Ocampo-Peñuela et al., 2022). We retained species with an extent of occurrence in Colombia of at least 5,000 km², of which at least 10% fell within SINAP (Linero-Triana et al., 2023) (Appendix S1). We downloaded Global Biodiversity Information Facility (GBIF) occurrence records from 2000 to 2022, kept those with coordinate uncertainty ≤1 km, and thinned them to 1 record per species per 5-km cell to reduce sampling bias (Steen et al., 2021). We retained only species with at least 20 records. The final data set comprised 181 species (141 birds and 40 mammals) with 287,563 bird and 10,767 mammal records (Appendix S2). It included forest-interior species and species of savannas, shrublands, and modified landscapes.

We characterized each species by its exposure to human modification and by traits related to movement. Exposure was the median change in the global Human Modification index (gHM; Theobald et al., 2025) between 2000 and 2022 (ΔgHM) at the species’ records. We used change rather than a single-year value to capture the landscape trajectory each species experienced. Traits were body mass, dispersal distance, trophic level, diet breadth, habitat breadth, and clutch or litter size (Broekman et al., 2022; Soria et al., 2021; Tobias et al., 2022) (Appendix S2). We estimated bird dispersal from the hand-wing index (Ocampo-Peñuela et al., 2023). We obtained mammal dispersal distances from the COMBINE database (Soria et al., 2021), which estimates them with allometric models (Santini et al., 2013). We set a minimum dispersal distance of 0.9 km (3 pixels).

Following Prima et al. (2024), we built a weighted dissimilarity matrix that assigned 25% of the weight to dispersal distance, 25% to ΔgHM, and 10% to each of the remaining 5 traits. We used Euclidean distance for continuous traits and Gower distance for trophic level, and rescaled each distance matrix to 0–1 before weighting. We applied Ward.D2 hierarchical clustering separately to birds and mammals and compared the Dunn index and average silhouette width across 2–15 groups with the fpc package (Hennig, 2026) and retained 6 groups per class (Appendix S3).

### 2.4. Connectivity modeling

We modeled connectivity for each EFG with Omniscape v0.5.8 in Julia v1.11.6 (Landau et al., 2021). Omniscape requires 3 inputs: a source strength layer, which sets how much current each pixel can inject; a resistance surface, which represents how difficult it is to move through each pixel; and a search radius, the maximum distance over which a source can contribute current to a focal pixel (B. McRae et al., 2016). The algorithm moves a circular window across the landscape. At each position, it injects current from the source pixels inside the window and routes it through the resistance surface toward the focal pixel at the window center. Summing current across all windows yields a cumulative current map (Appendix S4)(Landau et al., 2021; B. McRae et al., 2016). Unlike least-cost approaches, this method represents movement along all possible pathways, including suboptimal ones, and its predictions agree with observed movement in validation studies (Dickson et al., 2019; McRae et al., 2008; Phillips et al., 2021). We set the search radius to the median dispersal distance of each EFG, so each window spans twice that distance. To minimize edge effects, we added a 56-km buffer around continental Colombia, equal to the largest dispersal distance among the 181 species (jaguar, *Panthera onca*) (Koen et al., 2014). Because each EFG includes species with different distributions, we modeled connectivity across the whole country, which represents the collective response of the group rather than the ranges of individual species (Prima et al., 2024; Sáenz-Jiménez et al., 2025).

We derived resistance from gHM for 2000 and 2022 with an exponential transformation (Keeley et al., 2016; Zeller et al., 2012):

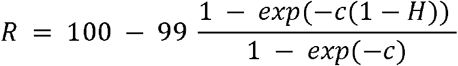

where *R* is resistance (1–100), *H* is gHM (0–1), and *c* controls the shape of the response. At low values of *c*, resistance increases almost linearly with modification (sensitive EFGs); at high values, resistance remains low until modification is high (tolerant EFGs) (Belote et al., 2022). We assigned the resistance sensitivity parameter c separately to each EFG, combining the median ΔgHM of its occurrence records with the documented sensitivity of its member species to habitat modification in the literature. This produced four levels: c = 0.25 (near-linear response, most sensitive groups) for AGF1, AGF2, MGF4 and MGF6; c = 1 for AGF5, AGF6, MGF1 and MGF3; c = 2 (moderate tolerance) for AGF4 and MGF5; and c = 4 (most tolerant groups) for AGF3 and MGF2 (Table S4.1). Low values make resistance rise steadily with human modification, whereas high values keep resistance low until modification is high (Keeley et al. 2016; Poor et al. 2025) (Appendix S4). Water bodies received specific resistance values so that they did not act as absolute barriers or current sinks (Clark et al., 2025; Poor et al., 2025). Source strength was the foreground area density (FAD) (Riitters & Wickham, 2012), of natural cover from MapBiomas Colombia Collection 3, resampled to 300 m for 2000 and 2022, so that more continuous natural cover injected more current. For each year, we calculated cumulated and normalized current (see above). We validated the 2022 models with GBIF records from 2018–2022 through the ratio of observed to expected records in areas of high normalized current (Creech et al., 2024) (Appendix S7).

### 2.5. Statistical matching

We used propensity score matching to reduce the selection bias inherent in PA designation, comparing protected pixels (treatment) with unprotected pixels of similar characteristics (control) (Andam et al., 2008; Schleicher et al., 2020). Following Negret et al. (2020), we matched each SIRAP independently. We estimated propensity scores with logistic regression, with elevation, slope, FAD in 2000, and gHM in 2000 as predictors of protection status (Appendix S6). We used gHM instead of the accessibility variables common in deforestation studies because it influences both where PAs were designated, since Colombia has favored areas of low modification (Correa-Ayram et al., 2020), and our response variable, since modification determines resistance to movement. We applied 1:1 nearest-neighbor matching without replacement, with exact matching by biome (IDEAM, 2024) and a caliper of 0.25 SD of the propensity score, in *MatchIt 4.7.2* (Ho et al., 2011). Treated pixels lay inside PAs declared through 2000. Control pixels lay outside any PA declared through 2022 and >5 km from PAs declared through 2000, to avoid spillover (Lessmann et al., 2024; Schleicher et al., 2020). We considered covariates balanced when the absolute standardized mean difference was <0.25. Each control inherited the PA of its matched treated pixel. We assigned each PA to the SIRAP containing most of its treated pixels and retained PAs with at least 10 matched pairs, which yielded 135 PAs. Spatial autocorrelation in the propensity score residuals was relevant (I > 0.1) only within 100 km, whereas treated pixels lay a median of 263 km from their matched controls (Appendix S6). Because matched pairs lay well beyond the range of this dependence, we considered that spatial autocorrelation did not bias the comparison between treated pixels and their controls.

### 2.6. Protection effects

We calculated 2 indicators of ecological multispecies connectivity change on the matched pixels, that is, the treated pixels of each PA and the control pixels matched to them. The first indicator, ΔZ, summarized all EFGs. For each EFG and SIRAP, we standardized the normalized current of each pixel as z scores:

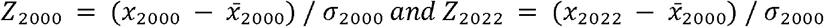

where x is the normalized current of a pixel, and x̅2022 and σ2000 are the mean and SD in 2000 across the treated and control pixels of the SIRAP (Bedson et al., 2025). Normalized current is cumulative current divided by the current expected iif resistance were uniform, so ΔZ reflects changes in permeability. Using the mean and SD of 2000 for both years placed them on the same scale. The change ΔZ = Z2022 – Z2000 expresses connectivity change in SD units of the matched pixels of each SIRAP. The multispecies ΔZ was the sum of the 12 EFG values, and the effect of each PA was the difference between the mean ΔZ of its treated pixels and that of the control pixels matched to them. We repeated the analysis with cumulative current.

The second indicator, CLT, also used the normalized current of each EFG. We classified pixels as impeded current (≤0.9), diffuse current (>0.9–1.3), or channelized current (>1.3) (Cameron et al., 2022; B. McRae et al., 2016; Prima et al., 2024). We defined critical loss as the transition of a pixel from a connected class (diffuse or channelized) in 2000 to impeded in 2022 (Fig. 2). For each EFG, we calculated CLT separately for the treated pixels of a PA and for its matched controls:

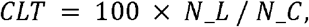

where N_C is the number of pixels with normalized current >0.9 in 2000 and N_L is the number of those pixels with values ≤0.9 in 2022. The multispecies CLT was the mean of the 12 EFG values.

Avoided CLT was the CLT of the controls minus that of the PA, in percentage points (pp), so positive values indicate that the PA lost less connectivity than its controls.

For both indicators, we tested whether PA effects differed from 0 with two-sided Wilcoxon signed-rank tests. We used two-sided tests because protection could increase as well as reduce connectivity loss (Negret et al., 2020). We estimated median effects with 95% confidence intervals (CIs) from 1000 percentile bootstrap iterations that resampled PAs. We report unadjusted p values because each test addressed a distinct question about a region, PA category, or functional group, and we interpret results through effect sizes and CIs rather than significance thresholds.

We analyzed all PAs together and disaggregated results by SIRAP, management category (strict protection, IUCN Ia–III; sustainable use, IUCN V), size (small, <1000 ha; medium, 1000 to <25,000 ha; large, ≥25,000 ha), and jurisdiction (national or regional) (Bonilla-Mejía & Higuera-Mendieta, 2019; Liévano-Latorre et al., 2021; Negret et al., 2020). We tested each category separately and did not compare categories formally, because management, size, and jurisdiction are partly confounded; differences among categories are therefore descriptive.

### 2.7. Magnitude and context of effects

To test whether the presence of an effect reflected its magnitude, we compared the mean CLT of the 12 EFGs in each PA with that of its controls. Control loss represents the loss expected without protection, that is, the pressure the PA faced (Pressey et al., 2015). The difference between control and PA loss is the avoided loss, in percentage points (pp), and its ratio to control loss is the proportion of avoided loss, in percent, which allows comparisons among PAs under different pressure.

We classified PAs into 5 categories. Protected areas that lost more than 1 pp less than their controls were effective, either under high pressure or under low or medium pressure. Those whose loss differed from that of their controls by 1 pp or less showed no effect, either despite pressure (control loss >1%) or because pressure was absent (control loss ≤1%). Those that lost more than 1 pp more than their controls formed the last category, greater loss than controls. Pressure levels corresponded to national terciles of control loss among PAs under pressure. The 1-pp margin is a management criterion rather than a statistical threshold.

## 3. Results

### 3.1. Emergent functional groups

We identified 12 EFGs, 6 of birds (AGF1–AGF6; 141 species) and 6 of mammals (MGF1–MGF6; 40 species), which differed in body mass, dispersal distance, and exposure to human modification (Table 1; Appendices S1 and S3).

**Table 1.**
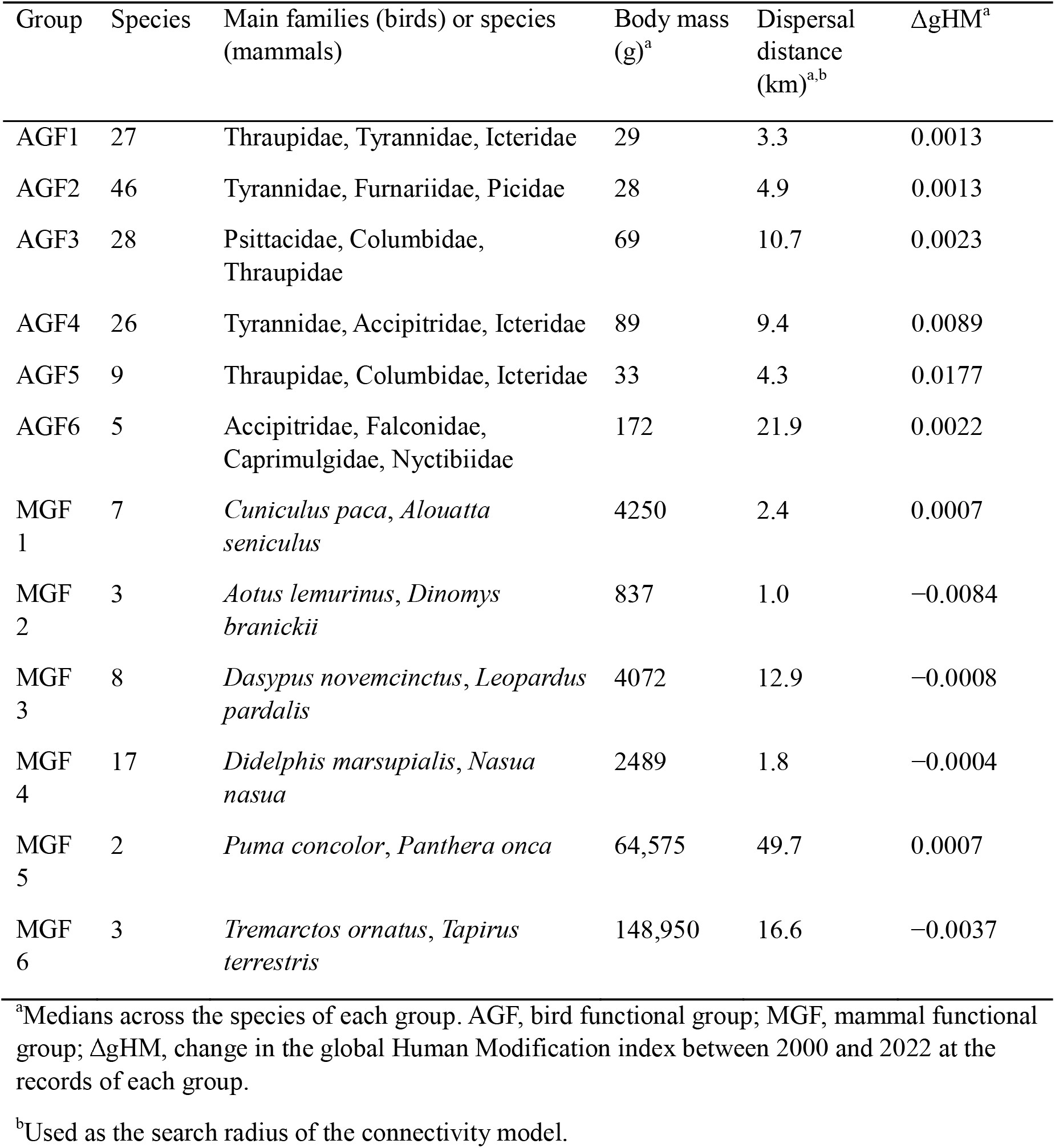
Number of species, main families or species, and median body mass, dispersal distance, and change in human modification of the 6 bird and 6 mammal emergent functional groups.

Among mammals, the 2 large felids, jaguar and puma (*Puma concolor*), formed MGF5, with the longest dispersal distance (median 49.7 km). Group MGF6 comprised the large herbivores and omnivores, lowland tapir (*Tapirus terrestris*), mountain tapir (*T. pinchaque*), and Andean bear (*Tremarctos ornatus*; median mass 149 kg; 16.6 km), and MGF3 carnivores and other species of intermediate mobility, such as the nine-banded armadillo (*Dasypus novemcinctus*), ocelot (*Leopardus pardalis*), and crab-eating raccoon (*Procyon cancrivorus*; 12.9 km). The other 3 groups had short dispersal distances: MGF1 comprised medium-sized herbivores and frugivores, such as the lowland paca (*Cuniculus paca*), red howler monkey (*Alouatta seniculus*), Central American agouti (*Dasyprocta punctata*), and brown-throated sloth (*Bradypus variegatus*; 2.4 km); MGF4, the largest mammal group (n = 17), mesocarnivores and omnivores, such as the common opossum (*Didelphis marsupialis*), crab-eating fox (*Cerdocyon thous*), tayra (*Eira barbara*), and South American coati (*Nasua nasua*; 1.8 km); and MGF2, 3 species with the shortest dispersal distance (1.0 km), lemurine night monkey (*Aotus lemurinus*), stump-tailed porcupine (*Coendou rufescens*), and pacarana (*Dinomys branickii*), recorded where human modification decreased the most (median change in gHM −0.008).

Among birds, AGF2, the largest group (n = 46), comprised insectivores and carnivores of the canopy and forest edge, mainly flycatchers (Tyrannidae, 13 species), ovenbirds (Furnariidae, 7), woodpeckers (Picidae, 6), and some raptors and cuckoos (median dispersal distance 4.9 km). Group AGF1 (n = 27) comprised small passerines, mainly tanagers (Thraupidae, 10 species), flycatchers (8), and orioles and blackbirds (Icteridae, 3), plus a few larger frugivores, such as the Wattled Guan (*Aburria aburri*; 3.3 km). Groups AGF3 (n = 28) and AGF5 (n = 9) comprised granivores and frugivores: AGF3 parrots (Psittacidae, 9 species), pigeons (Columbidae, 6), and tanagers (6; 10.7 km), and AGF5 tanagers, doves, and orioles, such as seedeaters (*Sporophila* spp.), Ruddy Ground Dove (*Columbina talpacoti*), and Yellow Oriole (*Icterus nigrogularis*; 4.3 km), which showed the largest increase in human modification (median change +0.018). Group AGF4 (n = 26) comprised generalist predators of open areas, hawks (Accipitridae, 5 species), falcons (Falconidae, 2), and cuckoos (Cuculidae, 2), together with flycatchers (6) and orioles and blackbirds (3; 9.4 km), and AGF6 (n = 5) diurnal raptors and nocturnal insectivores, such as the Bat Falcon (*Falco rufigularis*), Common Pauraque (*Nyctidromus albicollis*), and Common Potoo (*Nyctibius griseus*; 21.9 km).

### 3.2. Connectivity patterns

Connectivity patterns varied with the dispersal distance and resistance of each EFG (Appendix S5). Groups with short dispersal distances (MGF2, MGF4, and MGF1) produced fragmented flow, with high channelization restricted to short stretches of the Andean foothills, whereas EFGs with large search radii, such as the large felids (MGF5) and AGF6, produced more concentrated flow and better-defined corridors along the Eastern Cordillera, the Orinoco and Amazon foothills, and the continuous forest blocks of the Pacific.

Multispecies current in 2000 was highest along the Andean corridor, particularly in the Eastern Cordillera and its Orinoco and Amazon foothills, and in the large forest blocks of the Chocó and the northwestern Amazon, and lowest in the Caribbean and Orinoco plains and the north of the country. In 2022, current followed a similar pattern but had declined visibly in the Andes and increased slightly in parts of the Orinoco and its foothills. The most severe losses between 2000 and 2022 concentrated along the north–south Andean arc, from the Central Cordillera to the Colombian Massif, and mainly in the foothills of the Eastern Andes, within the Andes–Amazon Transition Belt (AATB), whereas the eastern Amazon and southern Pacific lost the least (Fig. 3).

**Figure 3.**
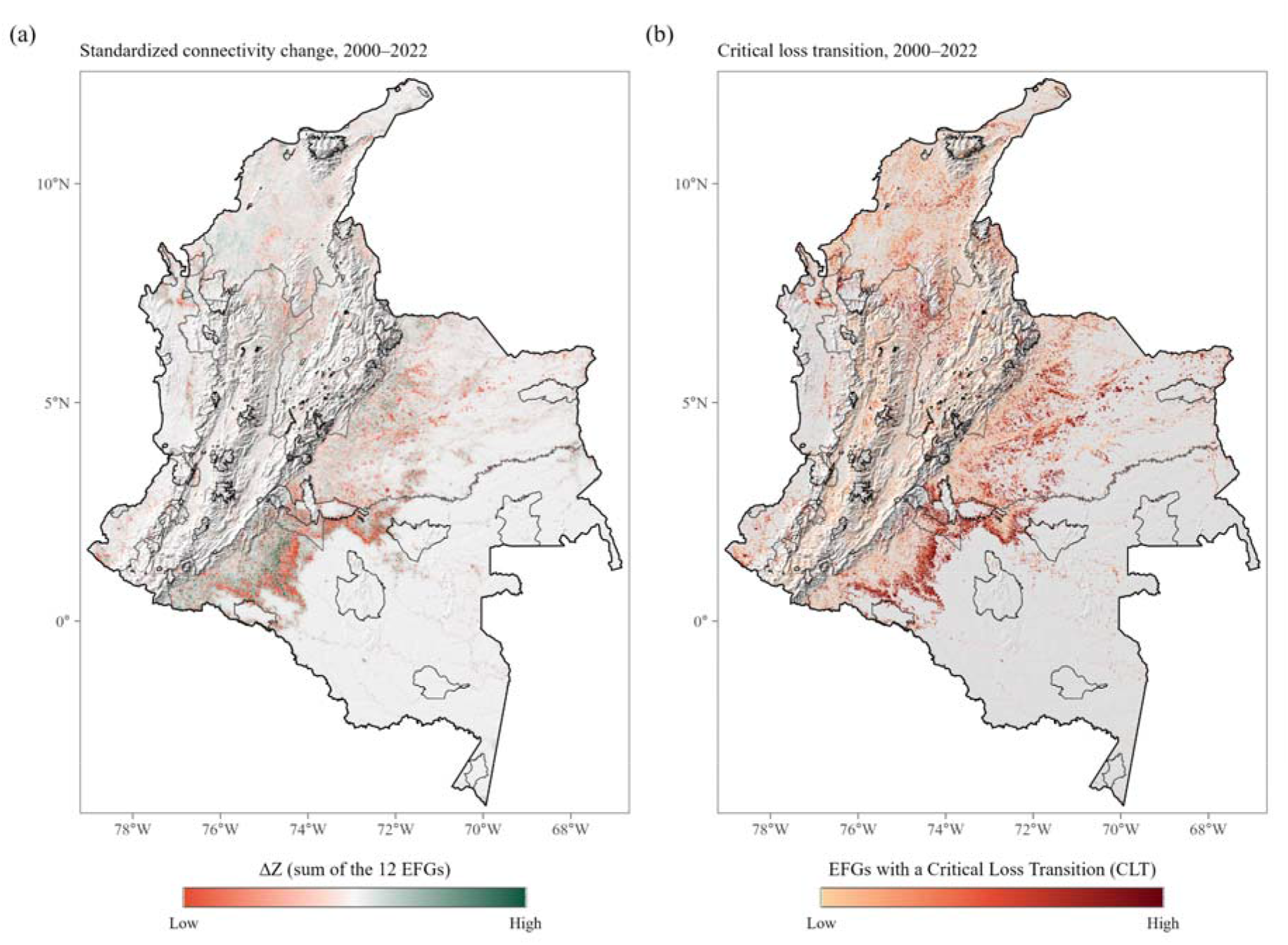
Change in multispecies functional connectivity in Colombia, 2000–2022. (a) Standardized connectivity change (ΔZ), summed across the 12 EFGs; red indicates loss and green indicates gain relative to 2000. (b) Critical loss transition (CLT), shown as the number of EFGs (out of 12) in which a pixel changed from connected in 2000 (normalized current >0.9) to impeded in 2022 (≤0.9). Thin black outlines show PAs declared by 2000.

### 3.3. Effects of protection

Matching retained 135 PAs and 937,143 treatment–control pairs in the 6 SIRAPs, with all covariates balanced (absolute standardized mean differences <0.1; Appendix S6).

Connectivity change (ΔZ) was more favorable in PAs than in their controls (median effect 0.87, 95% CI 0.36–1.60; p < 0.001), and 72.6% of PAs had positive effects. By SIRAP, effects were positive in the Caribbean, Northeastern Andes, Amazon, and Western Andes (p ≤ 0.018), tended to be positive in the Orinoco (p = 0.078), and were absent in the Pacific (Fig. 4; Table 2). Effects were positive in both management categories, in both jurisdictions, and in all size classes (p ≤ 0.040), and largest in small PAs (median 6.27; Fig. 5). With cumulative current, the national effect was similar (median 0.93, p < 0.001) and remained positive in the Amazon, Northeastern Andes, and Caribbean (p ≤ 0.013), but not in the Western Andes, Orinoco, or Pacific (p ≥ 0.12; Appendix S8).

**Figure 4.**
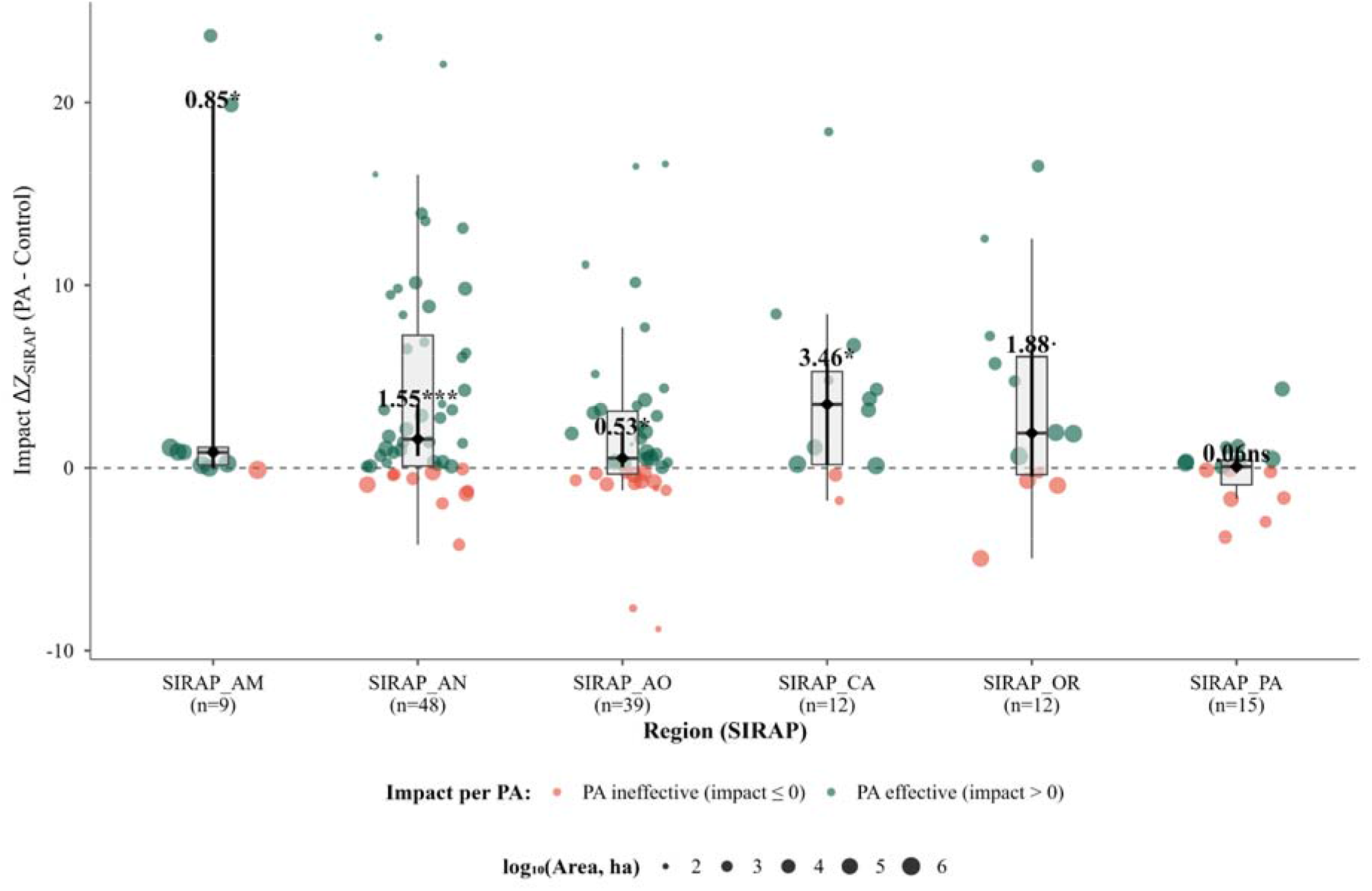
Effects of protection on standardized connectivity change (ΔZ; PA minus matched controls) for 135 PAs by regional subsystem (SIRAP). Each point is a PA, sized by the logarithm of its area (ha); points above 0 indicate PAs whose connectivity changed more favorably than that of their controls. Boxes show the median and interquartile range, lines show the median and its 95% bootstrap CI, and labels give the median with significance from two-sided Wilcoxon signed-rank tests (· p < 0.10, * p < 0.05, ** p < 0.01, *** p < 0.001; ns, not significant).

**Figure 5.**
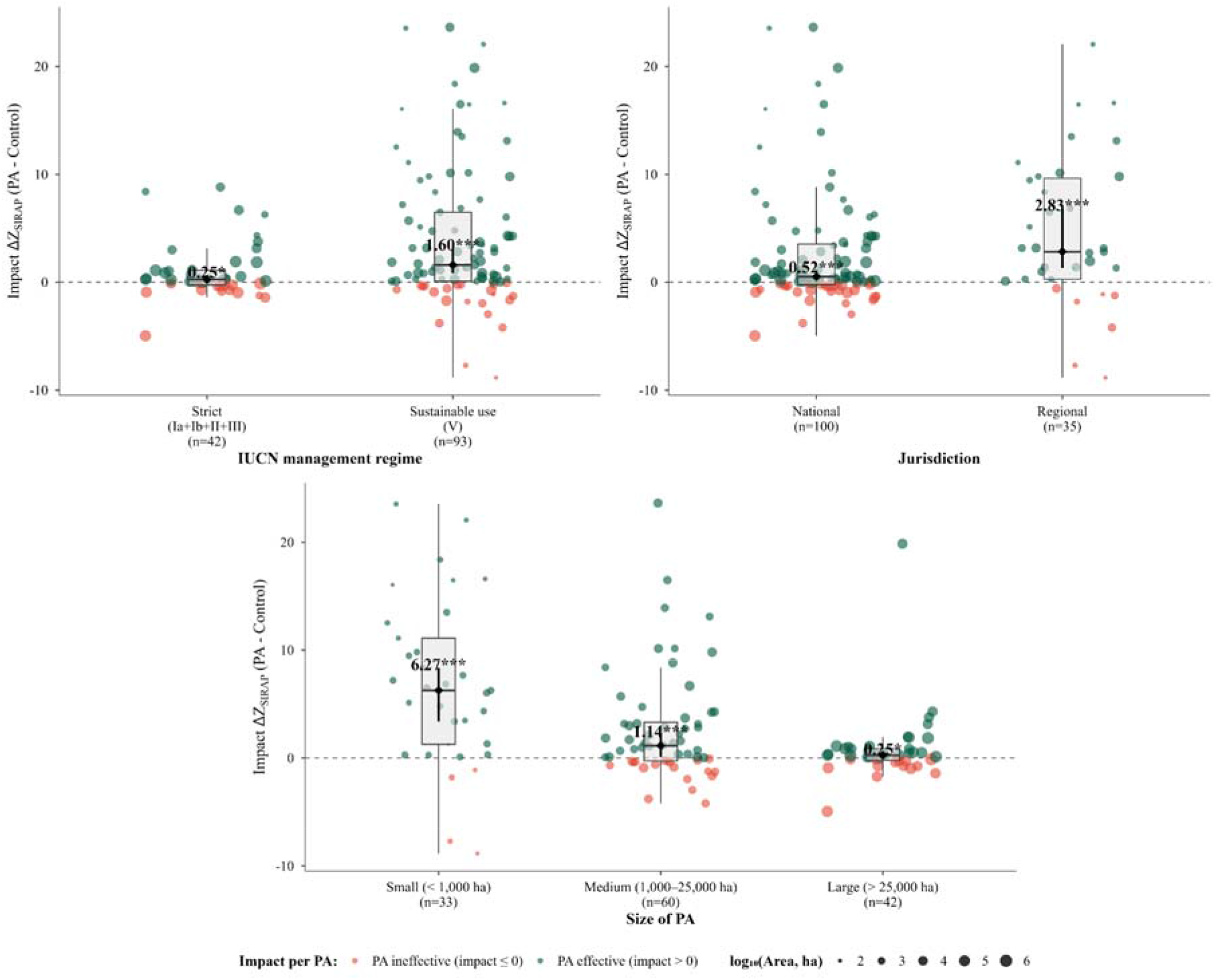
Effects of protection on standardized connectivity change (ΔZ; PA minus matched controls) by (a) management category (strict protection, International Union for Conservation of Nature categories Ia–III; sustainable use, category V), (b) jurisdiction (national or regional), and (c) size (small, <1000 ha; medium, 1000 to <25,000 ha; large, ≥25,000 ha). Symbols as in Fig. 4.

**Table 2.** Effects of protection on standardized connectivity change and on avoided critical loss transition in 135 protected areas, nationally and by regional subsystem.

| Region | PAs | Effect on $\Delta Z$ (95% CI)<sup>a</sup> | p | Avoided CLT, pp (95% CI)<sup>a</sup> | p |
| --- | --- | --- | --- | --- | --- |
| National | 135 | 0.87 (0.36 to 1.60) | <0.001 | 1.51 (0.38 to 2.74) | 0.003 |
| Amazon | 9 | 0.85 (0.003 to 19.9) | 0.018 | 0.38 (−0.92 to 19.7) | 0.477 |
| Northeastern Andes | 48 | 1.55 (0.66 to 3.49) | <0.001 | 3.41 (1.12 to 6.03) | 0.005 |
| Western Andes | 39 | 0.53 (0.04 to 1.95) | 0.010 | 0.64 (−0.97 to 2.73) | 0.630 |
| Caribbean | 12 | 3.46 (0.16 to 5.74) | 0.017 | 3.01 (0.30 to 7.41) | 0.055 |
| Orinoco | 12 | 1.88 (−0.48 to 6.46) | 0.078 | 5.47 (−1.71 to 14.6) | 0.147 |
| Pacific | 15 | 0.06 (−0.21 to 0.50) | 0.977 | −1.58 (−5.40 to 1.42) | 0.182 |

<sup>a</sup>Effects are medians of the differences between protected areas (PAs) and matched controls, with bootstrap 95% CIs. $\Delta Z$ , standardized connectivity change (SD units summed over 12 emergent functional groups); CLT, critical loss transition; pp, percentage points. Positive values indicate that PAs lost less connectivity than their controls. p values are from two-sided Wilcoxon signed-rank tests and are not adjusted.

Critical loss in matched controls, which approximates the loss expected without protection, varied among EFGs from 6.6% in AGF3 to 12.1% in AGF1, and was highest in AGF1, MGF1, and MGF3 (11.1–12.1%) and lowest in AGF3, MGF5, and AGF4 (6.6–8.0%). Protection reduced critical loss nationally (median avoided CLT 1.51 pp, 95% CI 0.38–2.74; p = 0.003). Protected areas lost less than their controls in all 12 EFGs, and the difference was significant for 8 (1.4–2.4 pp), with the largest values in MGF4, MGF5, and AGF6 (2.1–2.4 pp), but not for AGF1, AGF2, MGF1, and MGF3 (0.2–1.2 pp). By SIRAP, the multispecies effect was significant only in the Northeastern Andes (3.41 pp; p = 0.005), with a tendency in the Caribbean (3.01 pp; p = 0.055; Table 2). In the Northeastern Andes, PAs lost less than their controls in all 12 EFGs, significantly in 9 (5 bird and 4 mammal EFGs); in the Caribbean, significantly in 2 (AGF3 and AGF6), and in the other SIRAPs, in none (Fig. 6).

**Figure 6.**
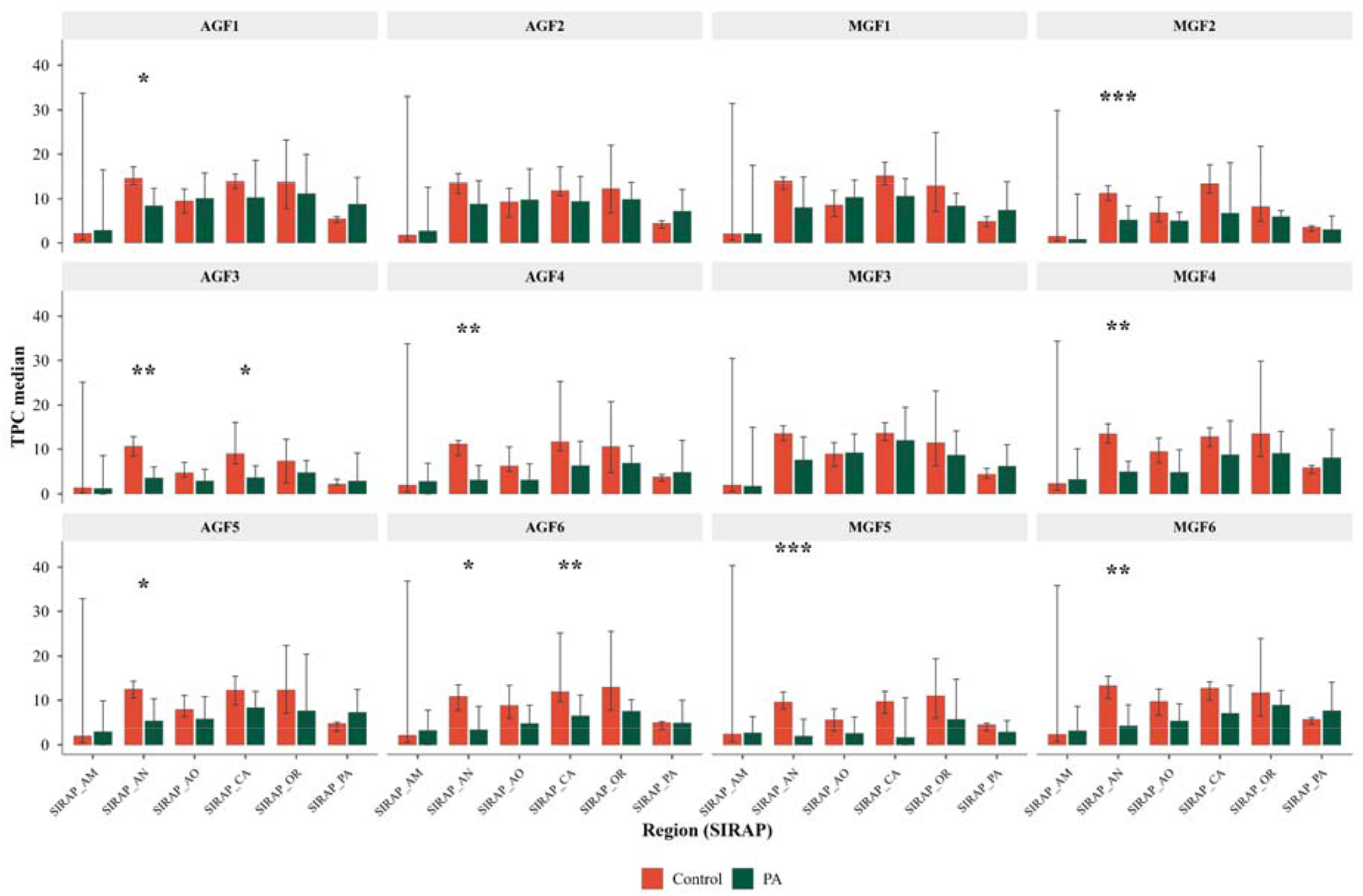
Median critical loss transition (CLT, %) in protected areas (PAs) and their matched controls for 6 bird (AGF1–AGF6) and 6 mammal (MGF1–MGF6) emergent functional groups by regional subsystem (SIRAP). Error bars are 95% bootstrap CIs of the median, and asterisks indicate significant differences between PAs and controls (two-sided Wilcoxon signed-rank tests on per-PA differences; * p < 0.05, ** p < 0.01, *** p < 0.001).

Relative to pressure, 74 of 135 PAs (55%) were effective, 33 under high pressure and 41 under low or medium pressure; 19 (14%) showed no effect despite pressure, 2 (1.5%) faced no pressure, and 40 (30%) lost more than their controls (median excess loss 5.9 pp; Fig. 7). Terciles of control loss were 6.1% and 13.4%. Sustainable-use PAs avoided a median of 2.6 pp, compared with 0.9 pp in strict-protection PAs, and their controls lost more (median 12.0% vs 4.9%); the median proportion of expected loss avoided was 28% and 19%. In the Pacific, 8 of 15 PAs (53%) lost more than their controls, although these controls lost little (median 4.2%).

**Figure 7.**
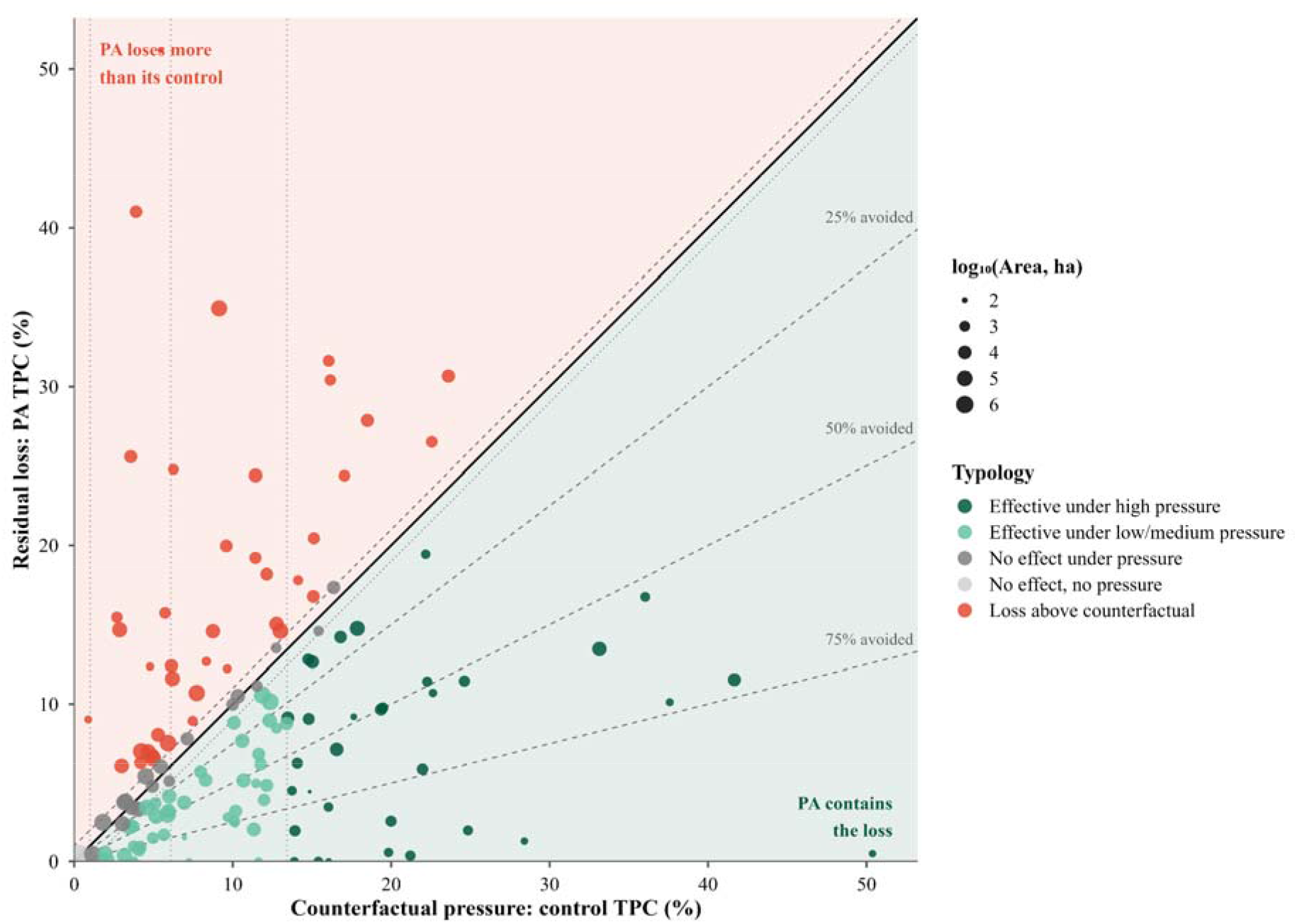
Critical loss transition (CLT, %) in each protected area (PA; y-axis, residual loss) relative to the CLT in its matched controls (x-axis, counterfactual pressure). Each point is a PA (n = 135), sized by the logarithm of its area (ha); CLT is the mean of the 12 emergent functional groups. The solid diagonal indicates equal loss; parallel dashed lines delimit the ±1 percentage point margin of no effect; dashed lines from the origin indicate 25%, 50%, and 75% of the expected loss avoided; and vertical dotted lines mark the absence of pressure (1%) and the national terciles of control loss (6.1% and 13.4%) that separate low, medium, and high pressure. Symbols identify the 5 effect categories (key on the figure).

## 4. Discussion

Protected areas maintained multispecies functional connectivity better than matched unprotected land in Colombia (hypothesis 1). To our knowledge, this is the first counterfactual evaluation of PAs that uses multispecies functional connectivity as the response variable. Between 2000 and 2022, 72.6% of PAs had more favorable connectivity trajectories than their matched controls, and nationally they avoided 1.5 pp of critical loss transitions (CLT; 95% CI 0.4–2.7). Effects on ΔZ were significant in 4 of the 6 SIRAPs, and PAs lost less critical connectivity than their controls in all 12 EFGs. The effects were clearest in the Northeastern Andes, where PAs avoided 3.4 pp of CLT, a median of 38% of the loss expected without protection, and largest in the Caribbean, where the effect on ΔZ reached 3.46 and PAs tended to avoid 3.0 pp of CLT. Grouping 181 species into 12 EFGs reduced the analysis dimensionality by 93% while retaining differences in how species move (Belote et al., 2022; Brennan et al., 2022; Prima et al., 2024). These results agree with evidence that Colombian PAs avoided deforestation (Bonilla-Mejía & Higuera-Mendieta, 2019; Liévano-Latorre et al., 2021; Negret et al., 2020) and extend it from forest cover to the movement of species.

The presence of an effect, however, overstates its contribution (hypothesis 3). The share of PAs with a favorable difference fell from 72.6% when counting any positive ΔZ to 63% when counting any avoided CLT, and to 55% when requiring more than 1 pp. How much loss an area can avoid depends on the pressure it faces (R. Pressey et al., 2015). A PA surrounded by land that lost little connectivity cannot avoid much, whereas a PA whose controls lost a great deal may avoid more in absolute terms but contain a smaller share of the expected loss (Forero-Medina et al., 2026).

Assessing only whether an effect exists confounds these scenarios, so we separated them into 5 categories. Of the 135 PAs, 74 were effective, 33 under high pressure and 41 under low or medium pressure; 2 had no effect because they faced no pressure, 19 had no effect despite pressure, and 40 lost more than their controls. Effectiveness rose with pressure, from 39% of PAs under low pressure to 55% under medium and 75% under high pressure (Joppa & Pfaff, 2011), partly because avoided loss cannot exceed control loss.

Effects varied among regions, in agreement with hypothesis 2, and followed the pressure each region faced. Avoided CLT was largest where controls lost the most (Northeastern Andes, Caribbean, and Orinoco; 3.0–5.5 pp). The Northeastern Andes showed the clearest effect, the only one significant on both indicators, with 9 of 12 EFGs benefiting. This region contained the most PAs (48 of 135) and the highest loss in matched controls (median 12.4%), so protection operated where there was most loss to avoid (Correa-Ayram et al., 2025). The Caribbean had the largest effect on ΔZ (3.46) and tended to avoid CLT, consistent with PAs that preserve remnants within the oldest and most transformed matrix of the country (Correa-Ayram et al., 2020). Los Colorados Fauna and Flora Sanctuary illustrates this role: it lost 9.8% of its critical connectivity, half the 19.5% lost by its controls. The Orinoco avoided the most loss (5.5 pp), but the interval was wide (n = 12 PAs) and the effect on ΔZ was only a tendency; gHM, which classifies grazing as modification, may overestimate resistance in natural savannas and reduce the contrast with controls (Poor et al., 2025; Theobald et al., 2025). This region also held the largest loss among strict-protection PAs: Tinigua National Natural Park lost 34.9%, compared with 9.1% in its controls, consistent with the deforestation documented in this park (Clerici et al., 2020; Murillo-Sandoval et al., 2022).

Where controls lost little, effects were negligible or negative (Western Andes, Pacific, and Amazon; −1.6 to 0.6 pp). In the Amazon, only 2 of 9 PAs faced high pressure, Serranía El Capricho and Serranía La Lindosa (controls lost 41.7% and 33.1%), and they avoided 30.1 and 19.7 pp; the other 7 had controls that lost 4.5% or less, which makes proportions unstable in this region. In the Western Andes, PAs lost as much as their controls (median 8.1% vs 8.0%), which suggests that they may have maintained flows without preventing barrier formation (Brennan et al., 2022). In the Pacific, controls lost a median of only 4.2%, reflecting a favorable baseline (Correa-Ayram et al., 2020; Vélez et al., 2020), yet 8 of 15 PAs lost more than their controls. The Andean-slope reserves Río Cali (41.0% vs 3.9% in its control) and La Elvira (25.6% vs 3.6%) stand out, whereas the neighboring Farallones de Cali National Natural Park lost 3.4% (4.6% in its control), which suggests that pressure originated within these reserves rather than in their surroundings (Clerici et al., 2020; González-González et al., 2021; Negret et al., 2020; Rubiano et al., 2026).

Effects also varied among functional groups (hypothesis 2), which supports evaluating connectivity for several groups rather than for one (Wood et al., 2022). Protection reduced critical loss in all 12 EFGs and significantly in 8 (1.4–2.4 pp), with the greatest effects in mesocarnivores and omnivores (MGF4), large felids (MGF5), and diurnal raptors (AGF6; 2.1–2.4 pp). A single focal species or a generic resistance surface would have hidden these differences(Koen et al., 2014; Wood et al., 2022). Dispersal distance alone did not explain the pattern. Effects were more frequent among groups that disperse farther (5 of 6 groups above 9 km) than among those that disperse less (3 of 6 below 5 km), yet MGF2 and MGF4, the groups with the shortest distances (1.0 and 1.8 km), benefited significantly, and MGF4 was among those that benefited most. Nor did a lack of pressure explain it: 3 of the 4 groups without a significant effect (AGF1, MGF1, and MGF3) had the highest losses in their controls, so they faced pressure and gained little. AGF1 consists of small generalist passerines that tolerate modified landscapes, and protection may alter their flow less than that of groups tied to continuous cover (Belote et al., 2022; Linero-Triana et al., 2023), but we did not test this explanation, and it does not apply to the other 3 groups. By region, 9 groups benefited in the Northeastern Andes and 2 in the Caribbean (AGF3 and AGF6), and elsewhere we found only tendencies.

Management category illustrates why magnitude matters (hypothesis 3)(Pressey et al., 2015) and highlights the role of sustainable-use PAs. Their effect on ΔZ was more than 6 times that of strict-protection PAs (median 1.60 vs 0.25). The advantage narrowed to 2.8 times for avoided CLT (2.6 vs 0.9 pp) and to 1.4 times for the proportion of expected loss avoided (28% vs 19%), because their controls also lost more (12.0% vs 4.9%), but it did not disappear. Sustainable-use PAs carried most of the protection where it was needed: they made up 39 of the 44 PAs under high pressure and 29 of the 33 PAs that were effective under it. Their outcomes were also more variable, as 34% lost more than their controls, compared with 19% of strict-protection PAs. These results align with studies showing that multiple-use PAs can be as effective as strict-protection PAs (Bonilla-Mejía & Higuera-Mendieta, 2019; Soares-Filho et al., 2010), a result that has been attributed to local governance by residents with formal rights (Alejo et al., 2021; Nelson & Chomitz, 2011) and to mosaics of lower resistance (Naidoo & Brennan, 2019). Because management is confounded with size, differences among categories remain descriptive: 76% of strict-protection PAs were large, whereas 89% of sustainable-use PAs were small or medium.

Size and jurisdiction followed the same logic as management, with one difference: the advantage of small PAs held in relative terms. Small PAs lie in the most transformed landscapes, where 58% faced high pressure, compared with 5% of large PAs, whereas large PAs tend to occupy remote areas with less loss to avoid (Joppa & Pfaff, 2009). Pressure alone, however, does not explain their performance, because they also contained the largest share of the loss expected without protection (median 70%, vs 8% in large PAs), consistent with evidence that small patches hold conservation value disproportionate to their area (Volenec & Dobson, 2020; Wintle et al., 2019). This advantage did not depend on the smallest samples, as it persisted after excluding PAs with fewer than 20 matched pixels (57%). Their effects were also the most variable, and part of their gains on ΔZ could reflect flow channeled toward remnants as the surrounding matrix loses permeability, rather than greater overall connectivity (McRae et al., 2008; Naidoo & Brennan, 2019). For this reason, we read their contribution through avoided CLT rather than ΔZ alone. Jurisdiction, in contrast, mattered little once pressure was considered: regional PAs avoided more critical loss than national PAs because their controls lost more, but both were equally effective under high pressure (75%). This suggests that regional environmental authorities achieve connectivity outcomes comparable to those of the national system when they face similar pressures.

These results do not settle the debate over a single large or several small reserves (SLOSS), because we compared effects per PA and not for equal total areas (Fahrig et al., 2022), but they add complementary evidence. Several small fragments can sustain as much biodiversity as a few large ones concerns species richness (Riva & Fahrig, 2022), whereas ours concerns connectivity: the 33 small PAs covered 0.13% of the area of the 135 PAs (12,604 of 9.9 million ha), yet the median small PA contained most of the loss expected without protection. This is compatible with a role as links or stepping stones within the network rather than as reservoirs of habitat. It contrasts with deforestation, for which large reserves were more effective (Liévano-Latorre et al., 2021), although those authors noted that small and regional PAs may serve other objectives, such as corridors that maintain ecological connectivity.

Our indicators complement structural metrics for Target 3 of the KM-GBF, which calls for well-connected systems of PAs and other effective area-based conservation measures (OECMs) (CBD, 2022). Structural indicators such as ProtConn describe how connected a system is, as the share of land that is protected and connected under a fixed dispersal threshold between polygons (10 km) (Saura et al., 2018, 2019). A system can meet such thresholds and still fail to prevent isolation driven by surrounding transformation (Belote et al., 2022; Brennan et al., 2022). Our indicators answer a different question: how much connectivity loss protection avoided, for 12 EFGs with distinct resistance and dispersal (median 1.0–49.7 km), over 22 years, and by region. Because they rest on a counterfactual comparison, they can be extended to OECMs, Indigenous reserves, Afro-Colombian collective territories, and peasant reserve zones by redefining the treatment and repeating the matching, without changing the connectivity surfaces. This extension is most needed in the Amazon and Pacific, where these territories cover large shares of the land (Anaya et al., 2020; Vélez et al., 2020).

For Colombia’s national protected area policy, our results support 3 directions. First, strengthen governance where protection did not translate into connectivity, as in the Andean-slope reserves of the Pacific and PNN Tinigua. Many PAs overlap or adjoin Indigenous reserves, Afro-Colombian collective territories, and peasant reserve zones, where environmental authority coexists with ethnic authorities and, in places, with illegal economies and armed actors (Anaya et al., 2020; Murillo-Sandoval et al., 2022; Vélez et al., 2020). In these contexts, coordination with ethnic authorities, joint management, institutional presence, and staff safety are preconditions for protection to translate into connectivity. Second, recognize the contribution of sustainable-use, regional, and small PAs, which carried most of the protection under high pressure, and conserve and restore the remnants that serve short-dispersal groups without a detectable effect (AGF1, AGF2, and MGF1), integrating them with networks already prioritized for wide-ranging species (Linero-Triana et al., 2023; Mosquera-Guerra et al., 2025; Zárrate-Charry et al., 2018). Third, build magnitude into how each PA is evaluated and managed: track the loss each area avoids relative to the pressure it faces, rather than the mere presence of an effect, and document the practices of PAs that remained effective under high pressure (Pressey et al., 2021). Beyond PA management, the indicators can provide a baseline for reporting the well-connected component of Target 3, a criterion for prioritizing new PAs, OECMs, and restoration under the national policy (DNP, 2021; MADS, 2024), and an input for the environmental guidelines that bind municipal land-use plans and for the assessment of linear infrastructure, where connectivity is still evaluated with structural criteria.

Our study has limitations that bear on how its effects should be read. Matching balances only the covariates we observed, so omitted territorial variables, such as overlap with Indigenous reserves and Afro-Colombian collective territories, armed conflict, local governance, and illegal economies, may bias the estimates; we therefore interpret them as conditional on the matched covariates (Lessmann et al., 2024; Schleicher et al., 2020). The connectivity surfaces rest on resistance derived from gHM, and omnidirectional models are sensitive to resistance and radius parameters, which may change the effects detected for some groups (Belote et al., 2022; Prima et al., 2024). We validated the models with occurrence records rather than movement data, so telemetry or capture– recapture data are the next step (Brennan et al., 2025; Creech et al., 2024). Two choices also shape the classification of effectiveness. The 1-pp margin is a management criterion: with margins of 0.5 and 2 pp, the share of effective PAs ranged from 59% to 47%, whereas effectiveness under high pressure barely changed (77% and 73%). Control loss, in turn, only approximates pressure.

Precision is lower for PAs with few pixels and for regions with few PAs (the Amazon, n = 9), and the 72 tests by EFG and SIRAP are exploratory and unadjusted. Effects on CLT clustered at the regional scale, so tests within each SIRAP are less affected by spatial dependence, whereas the national test, which treats PAs as independent, may overstate precision. Finally, we modeled only terrestrial connectivity, although hydrological connectivity matters for aquatic and semiaquatic fauna (Liczner et al., 2024), and we evaluated only SINAP PAs declared by 2000. Extending the matching, the validation, and the scope along these lines would test whether the contrast we found between the presence and the magnitude of effects holds.

Colombia’s protected areas maintained multispecies functional connectivity better than their counterfactuals, with effects in 4 of 6 regions. However, the presence of an effect overstated their contribution: only 55% of PAs avoided more than 1 pp of critical loss, and 30% lost more than their controls. Where pressure was high, protection worked in 3 of every 4 PAs, most of them sustainable-use areas. Measuring additionality relative to pressure offers a replicable framework, applicable in any country with land-cover maps, biodiversity records, and a delimited PA system, to assess whether PA systems are effectively well connected (CBD, 2022).

## 5. Data code and availability

Species occurrence records are available from GBIF (https://doi.org/10.15468/dl.ueyynx; https://doi.org/10.15468/dl.sufze5). The code used for species selection, connectivity modeling, matching, and effectiveness analyses will be archived in a public repository with a DOI upon publication.

## Supporting Information

Species prioritization and list (Appendix S1), occurrence records and functional traits (Appendix S2), definition of emergent functional groups (Appendix S3), connectivity modeling parameters (Appendix S4), connectivity outputs by functional group (Appendix S5), statistical matching and spatial autocorrelation (Appendix S6), validation of the connectivity models (Appendix S7).

## Notes

### Competing Interest Statement

The authors have declared no competing interest.

