## Supporting_Information for "Counterfactual effect of Colombian protected areas on multispecies functional connectivity"

**Multispecies functional connectivity: a tool to evaluate the effectiveness of Colombia’s protected areas**

The following Supporting Information is available online: Appendix S1 (species prioritization and species list), Appendix S2 (occurrence records and functional traits), Appendix S3 (definition of emergent functional groups), Appendix S4 (connectivity modeling parameters), Appendix S5 (connectivity outputs by emergent functional group), Appendix S6 (statistical matching and spatial autocorrelation), Appendix S7 (validation of connectivity models), and Appendix S8 (extended results).

### Appendix S1. Species prioritization and species list

#### Preliminary species list

In January 2026 we searched the Web of Science for recent (2015–2025) multispecies connectivity studies in Neotropical countries, using the query: ("Ecological connectivity" OR "Functional connectivity" OR "Landscape connectivity" OR "Corridors") AND ("Multispecies" OR "Multi-species" OR "Multiple species") AND ("Colombia" OR "Panama" OR "Venezuela" OR "Brazil" OR "Peru" OR "Ecuador" OR "Bolivia"). We complemented the results with gray literature from regional and national prioritization exercises. This produced a preliminary list of 350 species (260 birds and 90 mammals), mostly terrestrial. We standardized scientific names against the GBIF taxonomic backbone with the *taxize* R package (Chamberlain et al. 2026).

#### Prioritization criteria

We prioritized species with the greatest capacity to support connectivity at the national and regional scale (Linero-Triana et al. 2023) using two spatial-representativeness criteria applied to the potential-distribution models of Noguera-Urbano et al. (2023), which are maximum-entropy models at 1-km² resolution: (i) an extent of occurrence of at least 5,000 km² in Colombia and (ii) more than 10% of that extent inside the National System of Protected Areas (SINAP).

We excluded bats and aquatic or semi-aquatic mammals because robust dispersal information is lacking for the Neotropics. Among birds, we excluded aerial families (Apodidae, Trochilidae, Hirundinidae, Cathartidae), species whose primary habitat is wetland, artificial, marine, or desert, and species that are not resident according to BirdBase, because their high mobility and independence from terrestrial cover could overestimate connectivity (Ocampo-Peñuela et al. 2020). We also required a dispersal distance of at least 0.9 km (three 300-m pixels) and at least 20 records per species after thinning. The final list contains **181 species (141 birds and 40 mammals)** (Table S1.1). The list is not restricted to forest-interior species: it includes taxa associated with savannas, shrublands, and landscapes with some degree of human modification.

#### Species list

**Table S1.1.** Prioritized species (n = 181), assigned emergent functional group (EFG), and the traits used for clustering. Mass = body mass (g). Disp = dispersal distance (km). Clutch = clutch size (birds) or litter size (mammals). DB = diet breadth. HB = habitat breadth. TL = trophic level (1 = herbivore, 2 = omnivore, 3 = carnivore). HMI 2000 and HMI 2022 = median Human Modification Index of all records of the species in each year; ΔHMI = difference between the 2022 and 2000 medians (Theobald et al. 2025). Bird traits are from Tobias et al. (2022) and mammal traits from Soria et al. (2021).

| **#** | **EFG** | **Order** | **Family** | **Species** | **Mass** | **Disp** | **Clutch** | **DB** | **HB** | **TL** | **HMI 2000** | **HMI 2022** | **ΔHMI** |
| --- | --- | --- | --- | --- | --- | --- | --- | --- | --- | --- | --- | --- | --- |
| 1 | AGF3 | Tinamiformes | Tinamidae | *Crypturellus soui* | 216.2 | 9.87 | 2 | 3 | 4 | 1 | 0.093 | 0.104 | 0.000 |
| 2 | AGF1 | Galliformes | Cracidae | *Aburria aburri* | 1,405.1 | 2.75 | 2 | 2 | 2 | 1 | 0.039 | 0.047 | -0.002 |
| 3 | AGF1 | Galliformes | Odontophoridae | *Odontophorus hyperythrus* | 339.3 | 3.92 | 4 | 2 | 3 | 1 | 0.074 | 0.061 | -0.004 |
| 4 | AGF3 | Columbiformes | Columbidae | *Claravis pretiosa* | 68.2 | 10.82 | 2 | 2 | 3 | 1 | 0.120 | 0.138 | 0.002 |
| 5 | AGF5 | Columbiformes | Columbidae | *Columbina minuta* | 32.6 | 4.95 | 2 | 1 | 5 | 1 | 0.166 | 0.204 | 0.021 |
| 6 | AGF5 | Columbiformes | Columbidae | *Columbina talpacoti* | 46 | 5.83 | 2 | 1 | 5 | 1 | 0.178 | 0.203 | 0.008 |
| 7 | AGF3 | Columbiformes | Columbidae | *Leptotila verreauxi* | 146.9 | 7.44 | 2 | 3 | 5 | 1 | 0.148 | 0.160 | 0.001 |
| 8 | AGF3 | Columbiformes | Columbidae | *Patagioenas cayennensis* | 229 | 16.99 | 2 | 2 | 5 | 1 | 0.128 | 0.161 | 0.005 |
| 9 | AGF3 | Columbiformes | Columbidae | *Patagioenas goodsoni* | 176 | 16.33 | 2 | 4 | 1 | 1 | 0.023 | 0.030 | 0.001 |
| 10 | AGF3 | Columbiformes | Columbidae | *Patagioenas plumbea* | 178.8 | 16.84 | 1 | 2 | 2 | 1 | 0.051 | 0.063 | 0.001 |
| 11 | AGF3 | Columbiformes | Columbidae | *Zenaida auriculata* | 110.2 | 15.22 | 2 | 1 | 5 | 1 | 0.175 | 0.208 | 0.010 |
| 12 | AGF4 | Cuculiformes | Cuculidae | *Crotophaga ani* | 110.1 | 9.43 | 5 | 4 | 4 | 3 | 0.142 | 0.171 | 0.005 |
| 13 | AGF4 | Cuculiformes | Cuculidae | *Crotophaga major* | 148.3 | 8.62 | 5 | 4 | 6 | 2 | 0.170 | 0.198 | 0.011 |
| 14 | AGF2 | Cuculiformes | Cuculidae | *Crotophaga sulcirostris* | 82 | 5.39 | 4 | 3 | 3 | 3 | 0.230 | 0.246 | 0.005 |
| 15 | AGF2 | Cuculiformes | Cuculidae | *Dromococcyx phasianellus* | 84.5 | 10.09 | 4 | 2 | 3 | 3 | 0.033 | 0.047 | 0.001 |
| 16 | AGF2 | Cuculiformes | Cuculidae | *Tapera naevia* | 48.4 | 7.96 | 4 | 1 | 1 | 3 | 0.161 | 0.168 | 0.002 |
| 17 | AGF6 | Caprimulgiformes | Caprimulgidae | *Nyctidromus albicollis* | 57.8 | 17.50 | 2 | 1 | 4 | 3 | 0.125 | 0.146 | 0.002 |
| 18 | AGF6 | Nyctibiiformes | Nyctibiidae | *Nyctibius griseus* | 172 | 23.37 | 1 | 1 | 4 | 3 | 0.124 | 0.140 | 0.002 |
| 19 | AGF4 | Charadriiformes | Charadriidae | *Vanellus chilensis* | 327 | 13.98 | 4 | 2 | 5 | 3 | 0.162 | 0.197 | 0.011 |
| 20 | AGF4 | Pelecaniformes | Threskiornithidae | *Mesembrinibis cayennensis* | 756 | 9.57 | 3 | 3 | 5 | 3 | 0.108 | 0.157 | 0.010 |
| 21 | AGF2 | Accipitriformes | Accipitridae | *Accipiter bicolor* | 287.5 | 9.28 | 2 | 2 | 3 | 3 | 0.090 | 0.097 | 0.001 |
| 22 | AGF4 | Accipitriformes | Accipitridae | *Buteo nitidus* | 519 | 11.41 | 2 | 2 | 2 | 3 | 0.151 | 0.179 | 0.009 |
| 23 | AGF4 | Accipitriformes | Accipitridae | *Buteogallus meridionalis* | 808 | 14.05 | 1 | 2 | 6 | 3 | 0.165 | 0.214 | 0.015 |
| 24 | AGF2 | Accipitriformes | Accipitridae | *Chondrohierax uncinatus* | 286.1 | 6.64 | 2 | 1 | 5 | 3 | 0.146 | 0.163 | 0.003 |
| 25 | AGF4 | Accipitriformes | Accipitridae | *Gampsonyx swainsonii* | 92.9 | 18.01 | 2 | 2 | 4 | 3 | 0.237 | 0.278 | 0.022 |
| 26 | AGF4 | Accipitriformes | Accipitridae | *Geranospiza caerulescens* | 302.5 | 9.13 | 2 | 3 | 8 | 3 | 0.163 | 0.189 | 0.006 |
| 27 | AGF6 | Accipitriformes | Accipitridae | *Harpia harpyja* | 4,800 | 26.38 | 2 | 1 | 1 | 3 | 0.007 | 0.028 | 0.002 |
| 28 | AGF6 | Accipitriformes | Accipitridae | *Ictinia plumbea* | 249.9 | 21.90 | 2 | 1 | 2 | 3 | 0.088 | 0.123 | 0.006 |
| 29 | AGF4 | Accipitriformes | Accipitridae | *Rupornis magnirostris* | 269 | 11.78 | 2 | 2 | 5 | 3 | 0.132 | 0.151 | 0.002 |
| 30 | AGF2 | Accipitriformes | Accipitridae | *Spizaetus isidori* | 1,197.4 | 8.99 | 1 | 1 | 1 | 3 | 0.009 | 0.004 | -0.003 |
| 31 | AGF2 | Accipitriformes | Accipitridae | *Spizaetus ornatus* | 1,197.4 | 3.85 | 1 | 1 | 3 | 3 | 0.031 | 0.033 | -0.001 |
| 32 | AGF2 | Strigiformes | Strigidae | *Megascops choliba* | 132 | 6.64 | 2 | 2 | 3 | 3 | 0.142 | 0.159 | 0.004 |
| 33 | AGF4 | Coraciiformes | Alcedinidae | *Chloroceryle aenea* | 13.8 | 8.99 | 4 | 3 | 5 | 3 | 0.092 | 0.124 | 0.005 |
| 34 | AGF2 | Coraciiformes | Momotidae | *Momotus subrufescens* | 97.1 | 4.58 | 4 | 3 | 5 | 3 | 0.176 | 0.184 | 0.002 |
| 35 | AGF4 | Piciformes | Bucconidae | *Notharchus hyperrhynchus* | 95.9 | 10.53 | 2 | 3 | 4 | 3 | 0.032 | 0.048 | 0.002 |
| 36 | AGF2 | Piciformes | Bucconidae | *Nystalus radiatus* | 61 | 7.96 | 3 | 2 | 5 | 3 | 0.128 | 0.133 | 0.000 |
| 37 | AGF2 | Piciformes | Galbulidae | *Galbula ruficauda* | 26.5 | 6.71 | 3 | 1 | 7 | 3 | 0.166 | 0.174 | 0.002 |
| 38 | AGF2 | Piciformes | Picidae | *Campephilus melanoleucos* | 256 | 7.30 | 2 | 2 | 5 | 3 | 0.088 | 0.115 | 0.002 |
| 39 | AGF2 | Piciformes | Picidae | *Campephilus pollens* | 187.9 | 3.26 | 3 | 1 | 1 | 3 | 0.076 | 0.065 | -0.003 |
| 40 | AGF2 | Piciformes | Picidae | *Colaptes punctigula* | 62 | 4.88 | 3 | 1 | 6 | 3 | 0.179 | 0.204 | 0.007 |
| 41 | AGF4 | Piciformes | Picidae | *Melanerpes rubricapillus* | 52.3 | 9.43 | 4 | 2 | 5 | 2 | 0.191 | 0.203 | 0.004 |
| 42 | AGF2 | Piciformes | Picidae | *Picumnus granadensis* | 12.2 | 3.11 | 3 | 1 | 3 | 3 | 0.215 | 0.220 | -0.002 |
| 43 | AGF2 | Piciformes | Picidae | *Picumnus olivaceus* | 10.6 | 2.60 | 2 | 1 | 4 | 3 | 0.155 | 0.149 | 0.000 |
| 44 | AGF4 | Piciformes | Picidae | *Picumnus squamulatus* | 10.2 | 4.07 | 3 | 1 | 4 | 3 | 0.113 | 0.165 | 0.015 |
| 45 | AGF2 | Piciformes | Picidae | *Veniliornis kirkii* | 28.8 | 4.07 | 2 | 1 | 5 | 3 | 0.158 | 0.159 | 0.000 |
| 46 | AGF1 | Piciformes | Ramphastidae | *Ramphastos brevis* | 412 | 1.72 | 2 | 1 | 3 | 1 | 0.021 | 0.026 | 0.001 |
| 47 | AGF4 | Falconiformes | Falconidae | *Falco femoralis* | 335.8 | 23.96 | 3 | 2 | 4 | 3 | 0.187 | 0.238 | 0.021 |
| 48 | AGF6 | Falconiformes | Falconidae | *Falco rufigularis* | 163.6 | 20.80 | 3 | 2 | 1 | 3 | 0.121 | 0.144 | 0.002 |
| 49 | AGF2 | Falconiformes | Falconidae | *Herpetotheres cachinnans* | 623.6 | 5.10 | 2 | 1 | 4 | 3 | 0.125 | 0.147 | 0.002 |
| 50 | AGF2 | Falconiformes | Falconidae | *Micrastur semitorquatus* | 621.7 | 5.46 | 2 | 1 | 2 | 3 | 0.108 | 0.116 | 0.002 |
| 51 | AGF4 | Falconiformes | Falconidae | *Milvago chimachima* | 315.2 | 13.17 | 2 | 4 | 6 | 3 | 0.134 | 0.157 | 0.002 |
| 52 | AGF3 | Psittaciformes | Psittacidae | *Amazona ochrocephala* | 476.9 | 12.00 | 3 | 3 | 6 | 1 | 0.154 | 0.187 | 0.013 |
| 53 | AGF3 | Psittaciformes | Psittacidae | *Ara macao* | 1,015 | 15.22 | 2 | 4 | 4 | 1 | 0.063 | 0.115 | 0.008 |
| 54 | AGF3 | Psittaciformes | Psittacidae | *Ara severus* | 343 | 17.57 | 2 | 2 | 4 | 1 | 0.119 | 0.162 | 0.011 |
| 55 | AGF3 | Psittaciformes | Psittacidae | *Brotogeris jugularis* | 63.2 | 22.05 | 6 | 4 | 3 | 1 | 0.203 | 0.209 | 0.005 |
| 56 | AGF3 | Psittaciformes | Psittacidae | *Forpus conspicillatus* | 26.4 | 19.41 | 4 | 3 | 3 | 1 | 0.189 | 0.207 | 0.005 |
| 57 | AGF3 | Psittaciformes | Psittacidae | *Forpus passerinus* | 23 | 20.66 | 5 | 3 | 4 | 1 | 0.214 | 0.264 | 0.021 |
| 58 | AGF3 | Psittaciformes | Psittacidae | *Ognorhynchus icterotis* | 285 | 27.48 | 4 | 2 | 1 | 1 | 0.048 | 0.043 | -0.004 |
| 59 | AGF3 | Psittaciformes | Psittacidae | *Pionus menstruus* | 251 | 16.18 | 3 | 4 | 5 | 1 | 0.135 | 0.150 | 0.002 |
| 60 | AGF3 | Psittaciformes | Psittacidae | *Pyrrhura melanura* | 70.5 | 22.05 | 4 | 1 | 1 | 1 | 0.012 | 0.023 | 0.001 |
| 61 | AGF2 | Passeriformes | Cotingidae | *Pachyramphus cinnamomeus* | 20.3 | 6.42 | 4 | 2 | 5 | 3 | 0.122 | 0.131 | 0.000 |
| 62 | AGF4 | Passeriformes | Cotingidae | *Pachyramphus polychopterus* | 20.8 | 7.22 | 3 | 2 | 5 | 2 | 0.109 | 0.127 | 0.001 |
| 63 | AGF2 | Passeriformes | Cotingidae | *Pachyramphus rufus* | 18.8 | 6.49 | 4 | 2 | 4 | 3 | 0.170 | 0.173 | 0.000 |
| 64 | AGF3 | Passeriformes | Cotingidae | *Tityra inquisitor* | 43.1 | 10.67 | 2 | 2 | 2 | 1 | 0.137 | 0.152 | 0.004 |
| 65 | AGF3 | Passeriformes | Cotingidae | *Tityra semifasciata* | 79.3 | 9.43 | 2 | 3 | 4 | 1 | 0.075 | 0.084 | 0.000 |
| 66 | AGF3 | Passeriformes | Fringillidae | *Euphonia laniirostris* | 15 | 8.40 | 4 | 1 | 5 | 1 | 0.167 | 0.184 | 0.002 |
| 67 | AGF3 | Passeriformes | Fringillidae | *Euphonia trinitatis* | 11 | 8.69 | 4 | 2 | 5 | 1 | 0.227 | 0.249 | 0.011 |
| 68 | AGF2 | Passeriformes | Furnariidae | *Campylorhamphus trochilirostris* | 40.8 | 2.31 | 2 | 2 | 5 | 3 | 0.126 | 0.122 | -0.001 |
| 69 | AGF2 | Passeriformes | Furnariidae | *Dendrocincla fuliginosa* | 38.7 | 4.66 | 2 | 2 | 6 | 3 | 0.071 | 0.087 | 0.001 |
| 70 | AGF2 | Passeriformes | Furnariidae | *Furnarius leucopus* | 54.8 | 3.26 | 2 | 1 | 5 | 3 | 0.168 | 0.166 | -0.001 |
| 71 | AGF2 | Passeriformes | Furnariidae | *Lepidocolaptes souleyetii* | 25.7 | 6.71 | 2 | 2 | 4 | 3 | 0.161 | 0.164 | 0.000 |
| 72 | AGF2 | Passeriformes | Furnariidae | *Synallaxis albescens* | 11.2 | 1.35 | 3 | 1 | 5 | 3 | 0.140 | 0.152 | 0.001 |
| 73 | AGF2 | Passeriformes | Furnariidae | *Xenops minutus* | 10.6 | 5.32 | 2 | 1 | 2 | 3 | 0.022 | 0.033 | 0.000 |
| 74 | AGF2 | Passeriformes | Furnariidae | *Xiphorhynchus susurrans* | 46.7 | 4.29 | 2 | 2 | 6 | 3 | 0.174 | 0.173 | -0.001 |
| 75 | AGF4 | Passeriformes | Icteridae | *Cacicus cela* | 85.5 | 11.19 | 2 | 4 | 6 | 2 | 0.081 | 0.126 | 0.009 |
| 76 | AGF1 | Passeriformes | Icteridae | *Icterus auricapillus* | 32.2 | 6.93 | 2 | 3 | 3 | 2 | 0.153 | 0.167 | 0.003 |
| 77 | AGF1 | Passeriformes | Icteridae | *Icterus chrysater* | 53.4 | 4.73 | 2 | 3 | 4 | 2 | 0.153 | 0.165 | 0.000 |
| 78 | AGF5 | Passeriformes | Icteridae | *Icterus icterus* | 60.9 | 3.04 | 2 | 4 | 4 | 2 | 0.142 | 0.201 | 0.024 |
| 79 | AGF1 | Passeriformes | Icteridae | *Icterus mesomelas* | 39.9 | 1.94 | 3 | 3 | 5 | 2 | 0.149 | 0.161 | 0.001 |
| 80 | AGF5 | Passeriformes | Icteridae | *Icterus nigrogularis* | 40.2 | 5.10 | 2 | 3 | 7 | 2 | 0.216 | 0.242 | 0.014 |
| 81 | AGF4 | Passeriformes | Icteridae | *Molothrus bonariensis* | 41.5 | 11.26 | 3 | 2 | 5 | 2 | 0.196 | 0.223 | 0.011 |
| 82 | AGF4 | Passeriformes | Icteridae | *Psarocolius decumanus* | 206.3 | 9.43 | 2 | 4 | 5 | 2 | 0.107 | 0.134 | 0.003 |
| 83 | AGF4 | Passeriformes | Mimidae | *Mimus gilvus* | 52 | 7.96 | 3 | 2 | 3 | 3 | 0.149 | 0.191 | 0.013 |
| 84 | AGF2 | Passeriformes | Parulidae | *Myiothlypis fulvicauda* | 14.9 | 2.45 | 2 | 1 | 2 | 3 | 0.146 | 0.137 | -0.001 |
| 85 | AGF2 | Passeriformes | Parulidae | *Setophaga pitiayumi* | 6.8 | 5.98 | 4 | 3 | 4 | 3 | 0.145 | 0.147 | -0.001 |
| 86 | AGF1 | Passeriformes | Pipridae | *Manacus manacus* | 16.7 | 2.89 | 2 | 2 | 3 | 1 | 0.091 | 0.111 | 0.002 |
| 87 | AGF1 | Passeriformes | Thraupidae | *Coereba flaveola* | 10 | 5.32 | 3 | 3 | 5 | 1 | 0.149 | 0.166 | 0.003 |
| 88 | AGF1 | Passeriformes | Thraupidae | *Conirostrum leucogenys* | 7 | 7.44 | 2 | 2 | 3 | 2 | 0.225 | 0.229 | 0.001 |
| 89 | AGF3 | Passeriformes | Thraupidae | *Dacnis cayana* | 13 | 9.35 | 2 | 3 | 4 | 1 | 0.093 | 0.115 | 0.002 |
| 90 | AGF3 | Passeriformes | Thraupidae | *Dacnis lineata* | 11 | 10.01 | 2 | 2 | 3 | 1 | 0.096 | 0.102 | 0.000 |
| 91 | AGF2 | Passeriformes | Thraupidae | *Eucometis penicillata* | 27 | 4.58 | 2 | 2 | 6 | 3 | 0.126 | 0.122 | 0.000 |
| 92 | AGF2 | Passeriformes | Thraupidae | *Nemosia pileata* | 16 | 6.05 | 2 | 2 | 6 | 3 | 0.216 | 0.216 | 0.005 |
| 93 | AGF1 | Passeriformes | Thraupidae | *Ramphocelus dimidiatus* | 28 | 2.97 | 2 | 3 | 3 | 2 | 0.158 | 0.161 | 0.001 |
| 94 | AGF1 | Passeriformes | Thraupidae | *Saltator coerulescens* | 54.9 | 1.87 | 2 | 5 | 3 | 1 | 0.129 | 0.165 | 0.005 |
| 95 | AGF1 | Passeriformes | Thraupidae | *Saltator maximus* | 47.6 | 3.63 | 2 | 5 | 4 | 1 | 0.111 | 0.133 | 0.002 |
| 96 | AGF5 | Passeriformes | Thraupidae | *Saltator orenocensis* | 33.7 | 4.29 | 2 | 4 | 5 | 1 | 0.157 | 0.180 | 0.018 |
| 97 | AGF1 | Passeriformes | Thraupidae | *Saltator striatipectus* | 39 | 2.45 | 2 | 4 | 4 | 2 | 0.179 | 0.187 | 0.000 |
| 98 | AGF3 | Passeriformes | Thraupidae | *Sicalis flaveola* | 16.9 | 6.34 | 3 | 2 | 5 | 1 | 0.189 | 0.219 | 0.012 |
| 99 | AGF5 | Passeriformes | Thraupidae | *Sporophila bouvronides* | 9.1 | 6.78 | 2 | 1 | 2 | 1 | 0.128 | 0.148 | 0.018 |
| 100 | AGF5 | Passeriformes | Thraupidae | *Sporophila intermedia* | 12.1 | 3.33 | 2 | 3 | 6 | 1 | 0.185 | 0.210 | 0.009 |
| 101 | AGF5 | Passeriformes | Thraupidae | *Sporophila lineola* | 9.8 | 3.99 | 2 | 1 | 3 | 1 | 0.106 | 0.171 | 0.035 |
| 102 | AGF5 | Passeriformes | Thraupidae | *Sporophila minuta* | 7.8 | 4.29 | 2 | 1 | 4 | 1 | 0.173 | 0.199 | 0.007 |
| 103 | AGF1 | Passeriformes | Thraupidae | *Sporophila nigricollis* | 9.6 | 2.01 | 2 | 1 | 4 | 1 | 0.159 | 0.171 | 0.001 |
| 104 | AGF3 | Passeriformes | Thraupidae | *Sporophila schistacea* | 12.7 | 9.50 | 2 | 1 | 4 | 1 | 0.215 | 0.229 | 0.002 |
| 105 | AGF1 | Passeriformes | Thraupidae | *Tachyphonus rufus* | 34.4 | 3.11 | 2 | 3 | 3 | 2 | 0.132 | 0.137 | 0.000 |
| 106 | AGF3 | Passeriformes | Thraupidae | *Tangara gyrola* | 21 | 6.71 | 2 | 2 | 3 | 1 | 0.121 | 0.117 | -0.001 |
| 107 | AGF3 | Passeriformes | Thraupidae | *Tangara johannae* | 20 | 8.99 | 2 | 2 | 3 | 1 | 0.014 | 0.014 | 0.000 |
| 108 | AGF1 | Passeriformes | Thraupidae | *Tiaris olivaceus* | 8.5 | 5.10 | 3 | 4 | 4 | 1 | 0.152 | 0.153 | -0.002 |
| 109 | AGF1 | Passeriformes | Thraupidae | *Volatinia jacarina* | 9.9 | 1.57 | 2 | 3 | 3 | 1 | 0.157 | 0.175 | 0.003 |
| 110 | AGF1 | Passeriformes | Turdidae | *Catharus aurantiirostris* | 29.8 | 3.26 | 2 | 3 | 6 | 2 | 0.137 | 0.123 | -0.003 |
| 111 | AGF1 | Passeriformes | Turdidae | *Turdus ignobilis* | 63.8 | 5.46 | 2 | 2 | 5 | 2 | 0.156 | 0.177 | 0.004 |
| 112 | AGF3 | Passeriformes | Turdidae | *Turdus leucomelas* | 69.1 | 7.66 | 3 | 2 | 5 | 1 | 0.147 | 0.173 | 0.008 |
| 113 | AGF2 | Passeriformes | Tyrannidae | *Camptostoma obsoletum* | 8.1 | 3.33 | 2 | 2 | 5 | 3 | 0.159 | 0.185 | 0.004 |
| 114 | AGF2 | Passeriformes | Tyrannidae | *Cnemotriccus fuscatus* | 13.6 | 1.57 | 3 | 1 | 4 | 3 | 0.108 | 0.117 | 0.003 |
| 115 | AGF2 | Passeriformes | Tyrannidae | *Contopus cinereus* | 11.6 | 7.44 | 2 | 1 | 2 | 3 | 0.156 | 0.174 | 0.001 |
| 116 | AGF1 | Passeriformes | Tyrannidae | *Elaenia flavogaster* | 24.8 | 3.26 | 2 | 2 | 5 | 2 | 0.170 | 0.188 | 0.004 |
| 117 | AGF3 | Passeriformes | Tyrannidae | *Legatus leucophaius* | 22.2 | 9.94 | 3 | 2 | 4 | 1 | 0.137 | 0.159 | 0.005 |
| 118 | AGF2 | Passeriformes | Tyrannidae | *Leptopogon amaurocephalus* | 11.7 | 4.95 | 2 | 2 | 3 | 3 | 0.093 | 0.115 | 0.002 |
| 119 | AGF4 | Passeriformes | Tyrannidae | *Machetornis rixosa* | 29.6 | 4.43 | 4 | 1 | 5 | 3 | 0.212 | 0.244 | 0.023 |
| 120 | AGF4 | Passeriformes | Tyrannidae | *Megarynchus pitangua* | 69.9 | 8.33 | 3 | 3 | 4 | 3 | 0.139 | 0.167 | 0.008 |
| 121 | AGF1 | Passeriformes | Tyrannidae | *Mionectes oleagineus* | 11.2 | 3.77 | 2 | 2 | 1 | 1 | 0.105 | 0.112 | 0.001 |
| 122 | AGF4 | Passeriformes | Tyrannidae | *Myiarchus crinitus* | 32.1 | 6.93 | 6 | 2 | 6 | 2 | 0.179 | 0.198 | 0.002 |
| 123 | AGF1 | Passeriformes | Tyrannidae | *Myiarchus panamensis* | 31.7 | 3.85 | 2 | 2 | 4 | 2 | 0.166 | 0.182 | 0.002 |
| 124 | AGF2 | Passeriformes | Tyrannidae | *Myiarchus tuberculifer* | 17.7 | 5.02 | 4 | 2 | 4 | 3 | 0.102 | 0.113 | 0.001 |
| 125 | AGF4 | Passeriformes | Tyrannidae | *Myiarchus venezuelensis* | 29.8 | 4.07 | 3 | 2 | 2 | 2 | 0.170 | 0.197 | 0.017 |
| 126 | AGF2 | Passeriformes | Tyrannidae | *Myiodynastes maculatus* | 43.2 | 6.12 | 3 | 3 | 5 | 3 | 0.155 | 0.165 | 0.001 |
| 127 | AGF2 | Passeriformes | Tyrannidae | *Myiopagis gaimardii* | 12 | 4.73 | 2 | 2 | 3 | 3 | 0.086 | 0.104 | 0.003 |
| 128 | AGF1 | Passeriformes | Tyrannidae | *Myiopagis viridicata* | 11.5 | 5.17 | 2 | 3 | 4 | 2 | 0.183 | 0.177 | -0.001 |
| 129 | AGF2 | Passeriformes | Tyrannidae | *Myiozetetes cayanensis* | 25.9 | 2.53 | 3 | 2 | 3 | 3 | 0.149 | 0.171 | 0.004 |
| 130 | AGF1 | Passeriformes | Tyrannidae | *Myiozetetes granadensis* | 29.3 | 3.11 | 3 | 2 | 4 | 2 | 0.074 | 0.104 | 0.004 |
| 131 | AGF2 | Passeriformes | Tyrannidae | *Myiozetetes similis* | 28 | 3.26 | 3 | 2 | 5 | 3 | 0.153 | 0.180 | 0.008 |
| 132 | AGF1 | Passeriformes | Tyrannidae | *Phaeomyias murina* | 10 | 2.53 | 2 | 3 | 5 | 2 | 0.182 | 0.197 | 0.005 |
| 133 | AGF1 | Passeriformes | Tyrannidae | *Phyllomyias griseiceps* | 7.2 | 4.43 | 3 | 2 | 1 | 2 | 0.177 | 0.174 | -0.002 |
| 134 | AGF4 | Passeriformes | Tyrannidae | *Pitangus sulphuratus* | 62.9 | 4.51 | 4 | 4 | 7 | 3 | 0.164 | 0.193 | 0.008 |
| 135 | AGF2 | Passeriformes | Tyrannidae | *Todirostrum cinereum* | 6.3 | 2.23 | 2 | 2 | 4 | 3 | 0.161 | 0.180 | 0.003 |
| 136 | AGF2 | Passeriformes | Tyrannidae | *Todirostrum nigriceps* | 6.5 | 3.26 | 1 | 1 | 4 | 3 | 0.141 | 0.148 | 0.001 |
| 137 | AGF2 | Passeriformes | Tyrannidae | *Tolmomyias sulphurescens* | 14.3 | 2.75 | 2 | 1 | 5 | 3 | 0.159 | 0.170 | 0.002 |
| 138 | AGF1 | Passeriformes | Tyrannidae | *Tyrannulus elatus* | 7 | 5.32 | 2 | 2 | 4 | 2 | 0.126 | 0.143 | 0.003 |
| 139 | AGF2 | Passeriformes | Tyrannidae | *Tyrannus melancholicus* | 37.4 | 8.62 | 2 | 2 | 4 | 3 | 0.142 | 0.171 | 0.004 |
| 140 | AGF4 | Passeriformes | Tyrannidae | *Tyrannus savana* | 31.9 | 14.71 | 2 | 2 | 5 | 3 | 0.166 | 0.207 | 0.015 |
| 141 | AGF2 | Passeriformes | Vireonidae | *Cyclarhis gujanensis* | 28.8 | 2.09 | 2 | 3 | 4 | 3 | 0.163 | 0.181 | 0.004 |
| 142 | MGF4 | Didelphimorphia | Didelphidae | *Chironectes minimus* | 856 | 0.97 | 3.62 | 3 | 2 | 2 | 0.109 | 0.095 | 0.000 |
| 143 | MGF4 | Didelphimorphia | Didelphidae | *Didelphis marsupialis* | 1,149.9 | 1.15 | 6.2 | 4 | 2 | 2 | 0.122 | 0.134 | 0.001 |
| 144 | MGF4 | Didelphimorphia | Didelphidae | *Marmosa robinsoni* | 98.5 | 0.90 | 9.75 | 2 | 2 | 2 | 0.104 | 0.136 | 0.001 |
| 145 | MGF3 | Cingulata | Dasypodidae | *Cabassous centralis* | 3,810 | 12.52 | 1 | 1 | 3 | 3 | 0.079 | 0.097 | -0.002 |
| 146 | MGF3 | Cingulata | Dasypodidae | *Dasypus novemcinctus* | 3,949 | 12.74 | 4 | 1 | 5 | 3 | 0.072 | 0.078 | -0.001 |
| 147 | MGF1 | Pilosa | Bradypodidae | *Bradypus variegatus* | 3,963.2 | 2.26 | 1 | 1 | 2 | 1 | 0.137 | 0.159 | 0.002 |
| 148 | MGF1 | Pilosa | Megalonychidae | *Choloepus hoffmanni* | 4,250 | 2.35 | 1 | 2 | 1 | 1 | 0.100 | 0.111 | -0.003 |
| 149 | MGF3 | Pilosa | Myrmecophagidae | *Myrmecophaga tridactyla* | 29,531.8 | 34.15 | 1 | 1 | 5 | 3 | 0.063 | 0.116 | 0.009 |
| 150 | MGF3 | Pilosa | Myrmecophagidae | *Tamandua mexicana* | 4,194.3 | 13.12 | 1 | 1 | 2 | 3 | 0.128 | 0.149 | -0.001 |
| 151 | MGF1 | Lagomorpha | Leporidae | *Sylvilagus floridanus* | 1,178.6 | 1.16 | 5 | 1 | 9 | 1 | 0.166 | 0.162 | -0.002 |
| 152 | MGF1 | Rodentia | Cuniculidae | *Cuniculus paca* | 9,000 | 3.55 | 1.01 | 3 | 1 | 1 | 0.041 | 0.042 | 0.001 |
| 153 | MGF1 | Rodentia | Dasyproctidae | *Dasyprocta punctata* | 2,229.6 | 1.65 | 1.64 | 2 | 3 | 1 | 0.078 | 0.078 | -0.001 |
| 154 | MGF2 | Rodentia | Dinomyidae | *Dinomys branickii* | 12,500 | 4.25 | 1.97 | 2 | 2 | 1 | 0.068 | 0.031 | -0.009 |
| 155 | MGF2 | Rodentia | Erethizontidae | *Coendou rufescens* | 831.8 | 1.04 | 1.284 | 2 | 1 | 1 | 0.113 | 0.101 | -0.007 |
| 156 | MGF4 | Rodentia | Heteromyidae | *Heteromys anomalus* | 69.3 | 0.90 | 2.415 | 2 | 2 | 2 | 0.141 | 0.136 | -0.003 |
| 157 | MGF4 | Rodentia | Heteromyidae | *Heteromys australis* | 267.5 | 0.90 | 2.75 | 2 | 1 | 2 | 0.014 | 0.028 | -0.002 |
| 158 | MGF2 | Primates | Aotidae | *Aotus lemurinus* | 837 | 0.96 | 1 | 4 | 1 | 2 | 0.071 | 0.057 | -0.008 |
| 159 | MGF1 | Primates | Atelidae | *Alouatta seniculus* | 6,398.3 | 2.94 | 1.2 | 2 | 2 | 1 | 0.098 | 0.134 | 0.001 |
| 160 | MGF1 | Primates | Atelidae | *Ateles hybridus* | 6,692.4 | 3.02 | 1 | 1 | 1 | 1 | 0.016 | 0.028 | 0.006 |
| 161 | MGF4 | Primates | Atelidae | *Lagothrix lagothricha* | 6,263.7 | 2.91 | 1 | 1 | 2 | 2 | 0.005 | 0.002 | 0.000 |
| 162 | MGF4 | Primates | Cebidae | *Cebus albifrons* | 2,488.6 | 1.75 | 1 | 4 | 2 | 2 | 0.050 | 0.074 | 0.001 |
| 163 | MGF4 | Primates | Cebidae | *Cebus capucinus* | 2,655 | 1.81 | 1 | 4 | 2 | 2 | 0.039 | 0.046 | -0.001 |
| 164 | MGF4 | Primates | Pitheciidae | *Pithecia milleri* | 1,537.5 | 1.56 | 1 | 3 | 1 | 2 | 0.024 | 0.009 | -0.001 |
| 165 | MGF4 | Carnivora | Canidae | *Cerdocyon thous* | 5,741.7 | 2.77 | 4 | 2 | 6 | 2 | 0.123 | 0.155 | 0.002 |
| 166 | MGF4 | Carnivora | Canidae | *Urocyon cinereoargenteus* | 4,220 | 2.34 | 3.895 | 4 | 2 | 2 | 0.056 | 0.069 | 0.002 |
| 167 | MGF3 | Carnivora | Felidae | *Leopardus pardalis* | 10,500 | 20.57 | 1.66 | 2 | 3 | 3 | 0.045 | 0.052 | 0.000 |
| 168 | MGF3 | Carnivora | Felidae | *Leopardus tigrinus* | 2,250 | 9.67 | 1.5 | 1 | 2 | 3 | 0.026 | 0.011 | -0.005 |
| 169 | MGF4 | Carnivora | Felidae | *Leopardus wiedii* | 3,600 | 2.14 | 1.5 | 2 | 3 | 2 | 0.032 | 0.026 | -0.002 |
| 170 | MGF5 | Carnivora | Felidae | *Panthera onca* | 81,150 | 56.04 | 2 | 1 | 6 | 3 | 0.034 | 0.039 | 0.001 |
| 171 | MGF5 | Carnivora | Felidae | *Puma concolor* | 48,000 | 43.32 | 2.76 | 1 | 5 | 3 | 0.042 | 0.046 | 0.001 |
| 172 | MGF3 | Carnivora | Mephitidae | *Conepatus semistriatus* | 1,598.7 | 8.18 | 3.995 | 1 | 3 | 3 | 0.131 | 0.177 | 0.008 |
| 173 | MGF4 | Carnivora | Mustelidae | *Eira barbara* | 4,135 | 2.31 | 2 | 1 | 2 | 2 | 0.048 | 0.051 | -0.002 |
| 174 | MGF4 | Carnivora | Mustelidae | *Galictis vittata* | 2,325 | 1.69 | 2 | 4 | 2 | 2 | 0.151 | 0.169 | 0.011 |
| 175 | MGF4 | Carnivora | Procyonidae | *Nasua nasua* | 4,000 | 2.27 | 3.81 | 2 | 2 | 2 | 0.023 | 0.017 | -0.003 |
| 176 | MGF4 | Carnivora | Procyonidae | *Potos flavus* | 2,420.9 | 1.72 | 1 | 1 | 1 | 2 | 0.067 | 0.067 | -0.002 |
| 177 | MGF3 | Carnivora | Procyonidae | *Procyon cancrivorus* | 6,270 | 15.98 | 2.75 | 2 | 2 | 3 | 0.131 | 0.142 | -0.001 |
| 178 | MGF6 | Carnivora | Ursidae | *Tremarctos ornatus* | 109,250 | 14.01 | 1.5 | 2 | 3 | 1 | 0.011 | 0.004 | -0.004 |
| 179 | MGF4 | Artiodactyla | Tayassuidae | *Tayassu pecari* | 31,798.7 | 7.11 | 2 | 3 | 4 | 2 | 0.020 | 0.026 | 0.001 |
| 180 | MGF6 | Perissodactyla | Tapiridae | *Tapirus pinchaque* | 148,950 | 16.61 | 1 | 1 | 4 | 1 | 0.005 | 0.002 | -0.004 |
| 181 | MGF6 | Perissodactyla | Tapiridae | *Tapirus terrestris* | 225,000 | 20.84 | 1 | 1 | 5 | 1 | 0.037 | 0.040 | 0.001 |

### Appendix S2. Occurrence records and functional traits

#### Occurrence records

We downloaded GBIF presence records for 2000–2022 (birds: GBIF.org, 24 February 2026, GBIF Occurrence Download, https://doi.org/10.15468/dl.ueyynx; mammals: GBIF.org, 20 February 2026, GBIF Occurrence Download, https://doi.org/10.15468/dl.sufze5). We kept only records with geographic coordinates, a documented basis of record, and a coordinate uncertainty of 1,000 m or less. We then spatially thinned the records on a 5-km grid with the *sf* and *terra* packages in R to reduce oversampling bias and ensure statistical independence (Steen et al. 2021). The final database contains 320,719 records for birds and 10,947 for mammals.

#### Functional traits and anthropogenic exposure

We characterized each species by (i) its exposure to anthropogenic change and (ii) six functional traits related to movement and connectivity (Table S2.1). Exposure is the median change in the Human Modification Index between 2000 and 2022 (ΔHMI; Theobald et al. 2025) within 5-km-radius local landscapes around each record (Bogoni et al. 2024; Ocampo-Peñuela et al. 2022). We used the change, rather than the static value of one year, to capture the transformation trajectory each species experienced during the study period.

Where empirical dispersal data were unavailable, we estimated dispersal distance from the hand-wing index for birds (dispersal = 0.734 × HWI − 8.777 km, with a minimum of 0.5 km; Ocampo-Peñuela et al. 2023) and from the allometric models of Santini et al. (2013), as compiled in COMBINE, for mammals (Soria et al. 2021), with a minimum of 0.9 km. Bird traits come from AVONET (Tobias et al. 2022) and BirdBase; trophic level follows the AVONET classification (scavengers were treated as carnivores). Clutch size is the midpoint of the reported range or, when missing, the family mean among the candidate species. Mammal traits come from COMBINE (Soria et al. 2021) and mammal home-range data from Broekman et al. (2022).

**Table S2.1.** Functional traits compiled and their expected relationship with vulnerability to connectivity loss.

| **Trait** | **Relevance for connectivity** | **References** |
| --- | --- | --- |
| Body mass | Species with greater body mass are more vulnerable to connectivity loss. | Cuervo and Møller 2020; Hillaert et al. 2018; Suárez-Castro et al. 2026 |
| Dispersal ability | Greater dispersal ability is associated with lower vulnerability to connectivity loss. | Santini et al. 2013; Suárez-Castro et al. 2026; Tobias et al. 2022 |
| Trophic level | Higher trophic levels are more vulnerable to connectivity loss. | Cuervo and Møller 2020; Hillaert et al. 2018; Suárez-Castro et al. 2026 |
| Diet breadth | Species with broader diets are less vulnerable to connectivity loss. | Cuervo and Møller 2020; Hillaert et al. 2018 |
| Habitat breadth | Species with broader habitat use are less vulnerable to connectivity loss. | Cuervo and Møller 2020; Hillaert et al. 2018 |
| Clutch size (birds) or litter size (mammals) | Species with greater reproductive potential are more tolerant of connectivity loss. | Cuervo and Møller 2020; Soria et al. 2021 |

### Appendix S3. Definition of emergent functional groups

We built a weighted dissimilarity matrix that assigned 25% to dispersal ability and 25% to ΔHMI, with the remaining 50% distributed equally among body mass, trophic level, diet breadth, habitat breadth, and clutch or litter size. We used Euclidean distances for continuous variables and Gower distance for trophic level (Prima et al. 2024). We then applied Ward.D2 hierarchical clustering independently to birds and mammals, following Boulangeat et al. (2012).

We chose k = 6 groups per taxon by evaluating the Dunn index and the mean silhouette width simultaneously for k = 2 to 15 with the *NbClust* package (Charrad et al. 2014; Figure S3.1). In mammals, the silhouette reached its maximum at k = 3 (S = 0.39) but dropped sharply at k = 4, and both indices plateaued from k = 6 onward with variation below 5% up to k = 15, indicating that additional groups do not substantially improve the clustering. In birds, k = 6 coincided with the inflection point of the Dunn curve, before the local maximum at k = 8–9, while the silhouette stayed stable at intermediate values (S ≈ 0.23–0.27) across the range. In both taxa, k < 6 produced groups that were too heterogeneous in dispersal ability and anthropogenic exposure, whereas k > 6 produced redundant groups with no interpretive gain. We therefore adopted k = 6 as a compromise between statistical parsimony, ecological coherence, and computational feasibility for the Omniscape modeling.

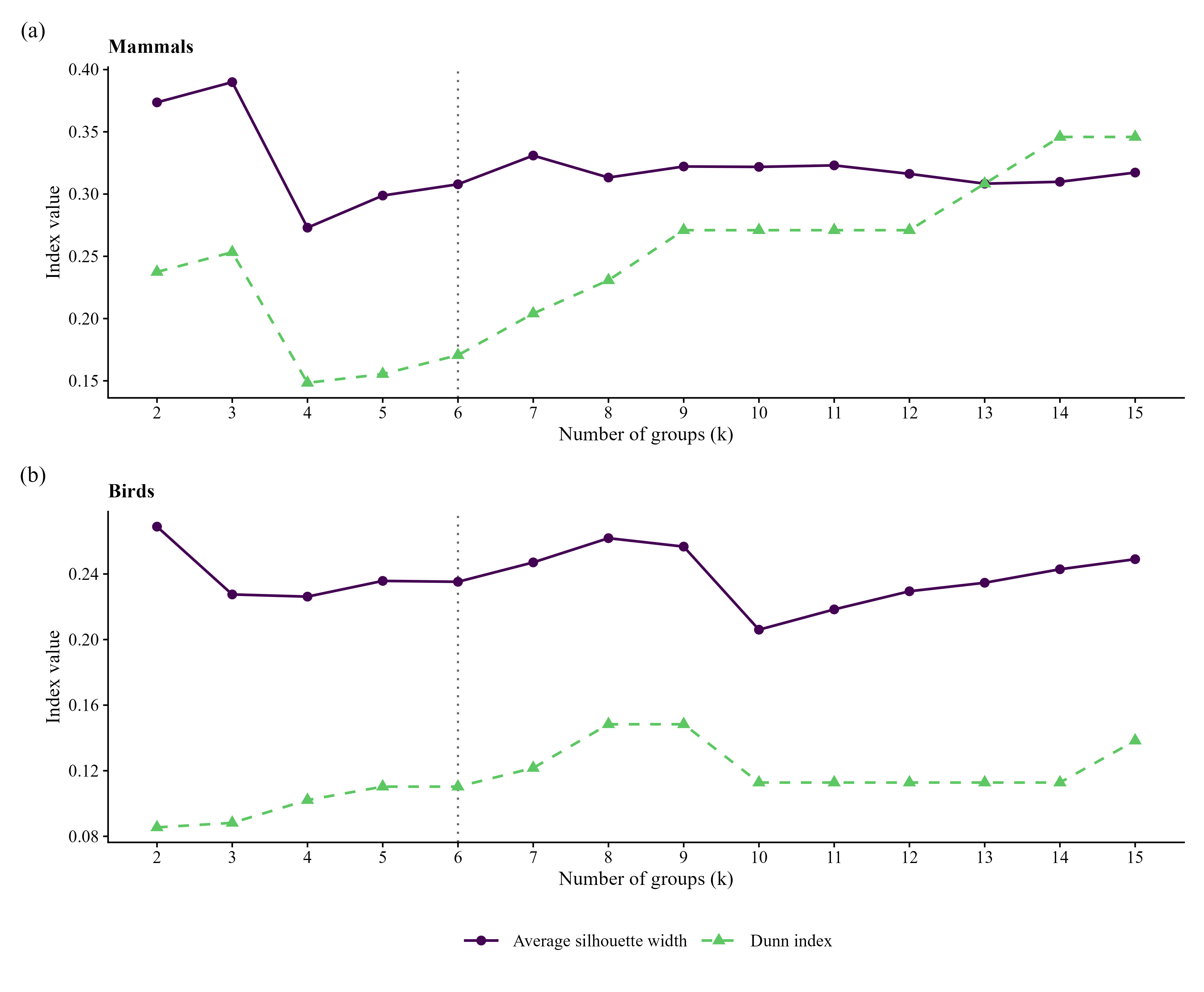

**Figure S3.1.** Dunn index and mean silhouette width for k = 2–15 groups, for mammals (top) and birds (bottom). The dashed vertical line marks the selected k = 6.

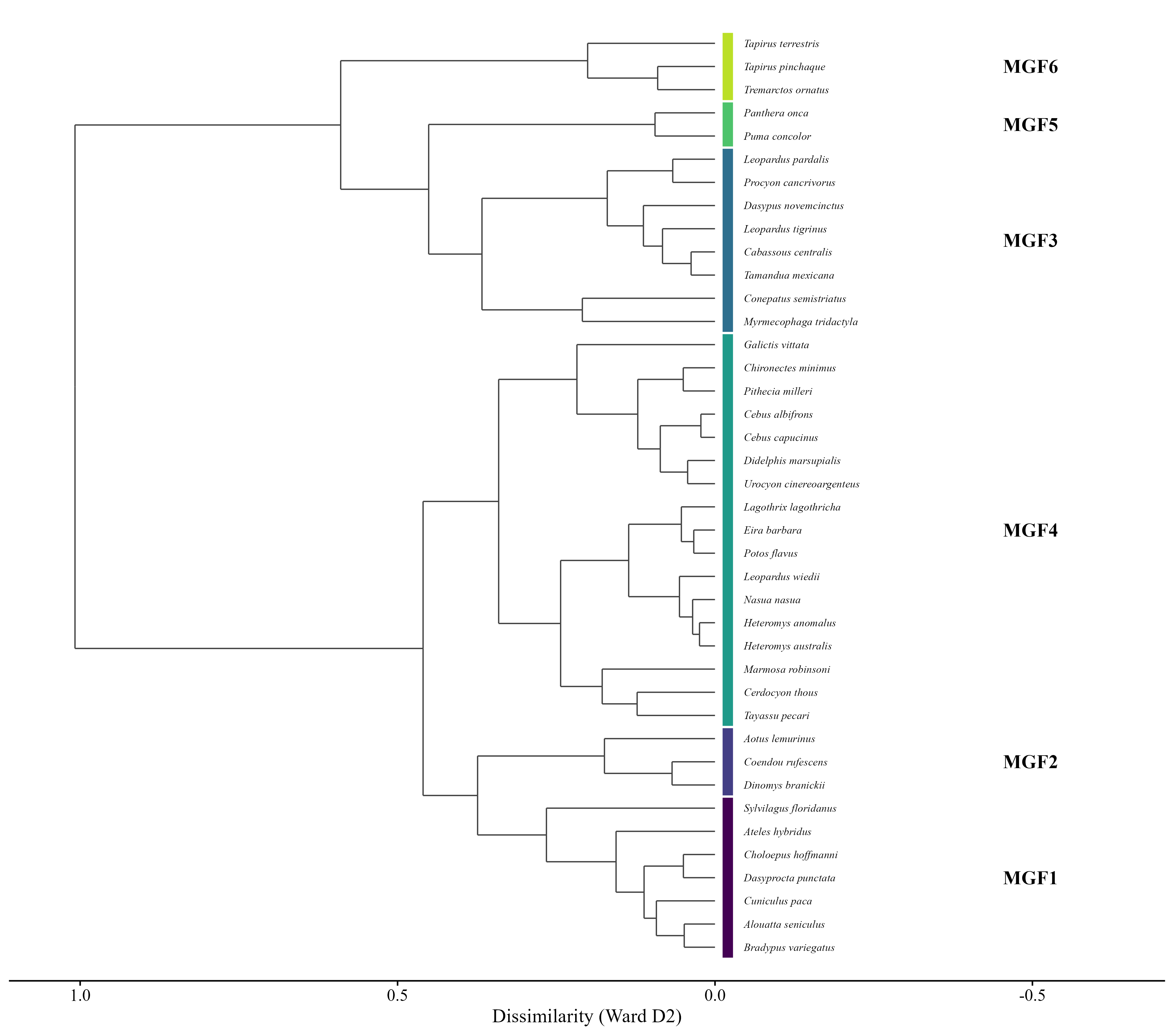

**Figure S3.2.** Dendrogram of the mammal emergent functional groups (EFGs; MGF1–MGF6) obtained by Ward D2 hierarchical clustering of life-history traits and anthropogenic exposure. Color identifies each EFG.

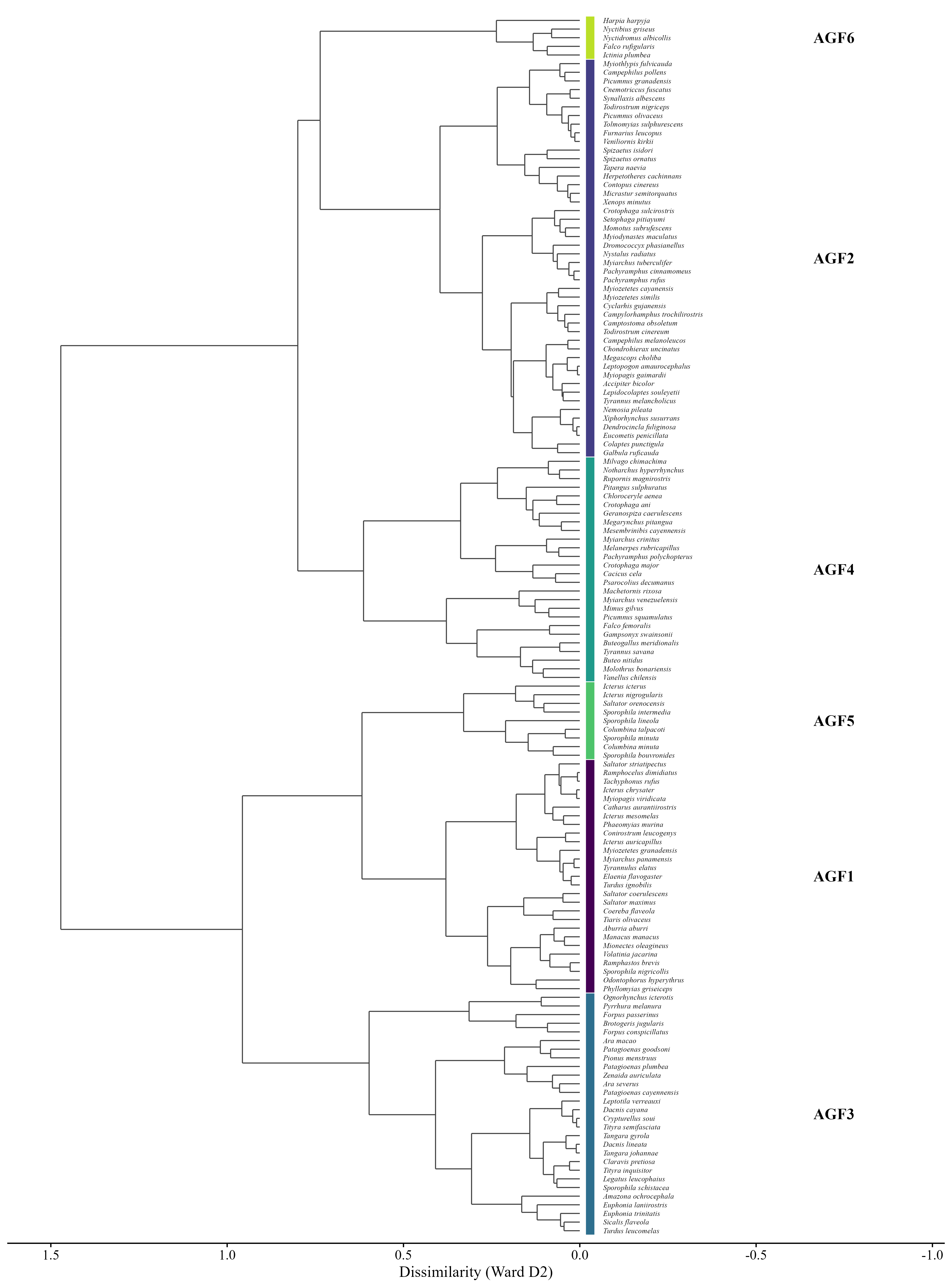

**Figure S3.3.** Dendrogram of the bird emergent functional groups (EFGs; AGF1–AGF6) obtained by Ward D2 hierarchical clustering of life-history traits and anthropogenic exposure. Color identifies each EFG.

**Table S3.1.** Traits of the emergent functional groups. Values are minimum–maximum (median), computed from Table S1.1. Dispersal medians in this table are the values used to set the Omniscape search radius (Appendix S4).

| **EFG** | **n** | **Body mass (g)** | **Dispersal (km)** | **ΔHMI** | **Clutch / litter** | **Diet breadth** | **Habitat breadth** | **Trophic level** |
| --- | --- | --- | --- | --- | --- | --- | --- | --- |
| AGF1 | 27 | 7.0–1,405.1 (29.3) | 1.57–7.44 (3.26) | -0.004–0.005 (0.001) | 2.00–4.00 (2.00) | 1.0–5.0 (3.0) | 1.0–6.0 (4.0) | omnivore 59%, herbivore 41% |
| AGF2 | 46 | 6.3–1,197.4 (28.4) | 1.35–10.09 (4.91) | -0.003–0.008 (0.001) | 1.00–4.00 (2.00) | 1.0–3.0 (2.0) | 1.0–7.0 (4.0) | carnivore 100% |
| AGF3 | 28 | 11.0–1,015.0 (68.7) | 6.34–27.48 (10.75) | -0.004–0.021 (0.002) | 1.00–6.00 (2.00) | 1.0–4.0 (2.0) | 1.0–6.0 (4.0) | herbivore 100% |
| AGF4 | 26 | 10.2–808.0 (89.2) | 4.07–23.96 (9.43) | 0.001–0.023 (0.009) | 1.00–6.00 (3.00) | 1.0–4.0 (2.0) | 2.0–8.0 (5.0) | carnivore 69%, omnivore 31% |
| AGF5 | 9 | 7.8–60.9 (32.6) | 3.04–6.78 (4.29) | 0.007–0.035 (0.018) | 2.00–2.00 (2.00) | 1.0–4.0 (1.0) | 2.0–7.0 (5.0) | herbivore 78%, omnivore 22% |
| AGF6 | 5 | 57.8–4,800.0 (172.0) | 17.50–26.38 (21.90) | 0.002–0.006 (0.002) | 1.00–3.00 (2.00) | 1.0–2.0 (1.0) | 1.0–4.0 (2.0) | carnivore 100% |
| MGF1 | 7 | 1,178.6–9,000.0 (4,250.0) | 1.16–3.55 (2.35) | -0.003–0.006 (0.001) | 1.00–5.00 (1.01) | 1.0–3.0 (2.0) | 1.0–9.0 (2.0) | herbivore 100% |
| MGF2 | 3 | 831.8–12,500.0 (837.0) | 0.96–4.25 (1.04) | -0.009–-0.007 (-0.008) | 1.00–1.97 (1.28) | 2.0–4.0 (2.0) | 1.0–2.0 (1.0) | herbivore 67%, omnivore 33% |
| MGF3 | 8 | 1,598.7–29,531.8 (4,071.6) | 8.18–34.15 (12.93) | -0.005–0.009 (-0.001) | 1.00–4.00 (1.58) | 1.0–2.0 (1.0) | 2.0–5.0 (3.0) | carnivore 100% |
| MGF4 | 17 | 69.3–31,798.7 (2,488.6) | 0.90–7.11 (1.75) | -0.003–0.011 (0.000) | 1.00–9.75 (2.00) | 1.0–4.0 (2.0) | 1.0–6.0 (2.0) | omnivore 100% |
| MGF5 | 2 | 48,000.0–81,150.0 (64,575.0) | 43.32–56.04 (49.68) | 0.001–0.001 (0.001) | 2.00–2.76 (2.38) | 1.0–1.0 (1.0) | 5.0–6.0 (5.5) | carnivore 100% |
| MGF6 | 3 | 109,250.0–225,000.0 (148,950.0) | 14.01–20.84 (16.61) | -0.004–0.001 (-0.004) | 1.00–1.50 (1.00) | 1.0–2.0 (1.0) | 3.0–5.0 (4.0) | herbivore 100% |

**Table S3.2.** Species composition of each emergent functional group.

| **EFG** | **Class** | **n** | **Species** |
| --- | --- | --- | --- |
| AGF1 | Birds | 27 | *Aburria aburri; Catharus aurantiirostris; Coereba flaveola; Conirostrum leucogenys; Elaenia flavogaster; Icterus auricapillus; Icterus chrysater; Icterus mesomelas; Manacus manacus; Mionectes oleagineus; Myiarchus panamensis; Myiopagis viridicata; Myiozetetes granadensis; Odontophorus hyperythrus; Phaeomyias murina; Phyllomyias griseiceps; Ramphastos brevis; Ramphocelus dimidiatus; Saltator coerulescens; Saltator maximus; Saltator striatipectus; Sporophila nigricollis; Tachyphonus rufus; Tiaris olivaceus; Turdus ignobilis; Tyrannulus elatus; Volatinia jacarina* |
| AGF2 | Birds | 46 | *Accipiter bicolor; Campephilus melanoleucos; Campephilus pollens; Camptostoma obsoletum; Campylorhamphus trochilirostris; Chondrohierax uncinatus; Cnemotriccus fuscatus; Colaptes punctigula; Contopus cinereus; Crotophaga sulcirostris; Cyclarhis gujanensis; Dendrocincla fuliginosa; Dromococcyx phasianellus; Eucometis penicillata; Furnarius leucopus; Galbula ruficauda; Herpetotheres cachinnans; Lepidocolaptes souleyetii; Leptopogon amaurocephalus; Megascops choliba; Micrastur semitorquatus; Momotus subrufescens; Myiarchus tuberculifer; Myiodynastes maculatus; Myiopagis gaimardii; Myiothlypis fulvicauda; Myiozetetes cayanensis; Myiozetetes similis; Nemosia pileata; Nystalus radiatus; Pachyramphus cinnamomeus; Pachyramphus rufus; Picumnus granadensis; Picumnus olivaceus; Setophaga pitiayumi; Spizaetus isidori; Spizaetus ornatus; Synallaxis albescens; Tapera naevia; Todirostrum cinereum; Todirostrum nigriceps; Tolmomyias sulphurescens; Tyrannus melancholicus; Veniliornis kirkii; Xenops minutus; Xiphorhynchus susurrans* |
| AGF3 | Birds | 28 | *Amazona ochrocephala; Ara macao; Ara severus; Brotogeris jugularis; Claravis pretiosa; Crypturellus soui; Dacnis cayana; Dacnis lineata; Euphonia laniirostris; Euphonia trinitatis; Forpus conspicillatus; Forpus passerinus; Legatus leucophaius; Leptotila verreauxi; Ognorhynchus icterotis; Patagioenas cayennensis; Patagioenas goodsoni; Patagioenas plumbea; Pionus menstruus; Pyrrhura melanura; Sicalis flaveola; Sporophila schistacea; Tangara gyrola; Tangara johannae; Tityra inquisitor; Tityra semifasciata; Turdus leucomelas; Zenaida auriculata* |
| AGF4 | Birds | 26 | *Buteo nitidus; Buteogallus meridionalis; Cacicus cela; Chloroceryle aenea; Crotophaga ani; Crotophaga major; Falco femoralis; Gampsonyx swainsonii; Geranospiza caerulescens; Machetornis rixosa; Megarynchus pitangua; Melanerpes rubricapillus; Mesembrinibis cayennensis; Milvago chimachima; Mimus gilvus; Molothrus bonariensis; Myiarchus crinitus; Myiarchus venezuelensis; Notharchus hyperrhynchus; Pachyramphus polychopterus; Picumnus squamulatus; Pitangus sulphuratus; Psarocolius decumanus; Rupornis magnirostris; Tyrannus savana; Vanellus chilensis* |
| AGF5 | Birds | 9 | *Columbina minuta; Columbina talpacoti; Icterus icterus; Icterus nigrogularis; Saltator orenocensis; Sporophila bouvronides; Sporophila intermedia; Sporophila lineola; Sporophila minuta* |
| AGF6 | Birds | 5 | *Falco rufigularis; Harpia harpyja; Ictinia plumbea; Nyctibius griseus; Nyctidromus albicollis* |
| MGF1 | Mammals | 7 | *Alouatta seniculus; Ateles hybridus; Bradypus variegatus; Choloepus hoffmanni; Cuniculus paca; Dasyprocta punctata; Sylvilagus floridanus* |
| MGF2 | Mammals | 3 | *Aotus lemurinus; Coendou rufescens; Dinomys branickii* |
| MGF3 | Mammals | 8 | *Cabassous centralis; Conepatus semistriatus; Dasypus novemcinctus; Leopardus pardalis; Leopardus tigrinus; Myrmecophaga tridactyla; Procyon cancrivorus; Tamandua mexicana* |
| MGF4 | Mammals | 17 | *Cebus albifrons; Cebus capucinus; Cerdocyon thous; Chironectes minimus; Didelphis marsupialis; Eira barbara; Galictis vittata; Heteromys anomalus; Heteromys australis; Lagothrix lagothricha; Leopardus wiedii; Marmosa robinsoni; Nasua nasua; Pithecia milleri; Potos flavus; Tayassu pecari; Urocyon cinereoargenteus* |
| MGF5 | Mammals | 2 | *Panthera onca; Puma concolor* |
| MGF6 | Mammals | 3 | *Tapirus pinchaque; Tapirus terrestris; Tremarctos ornatus* |

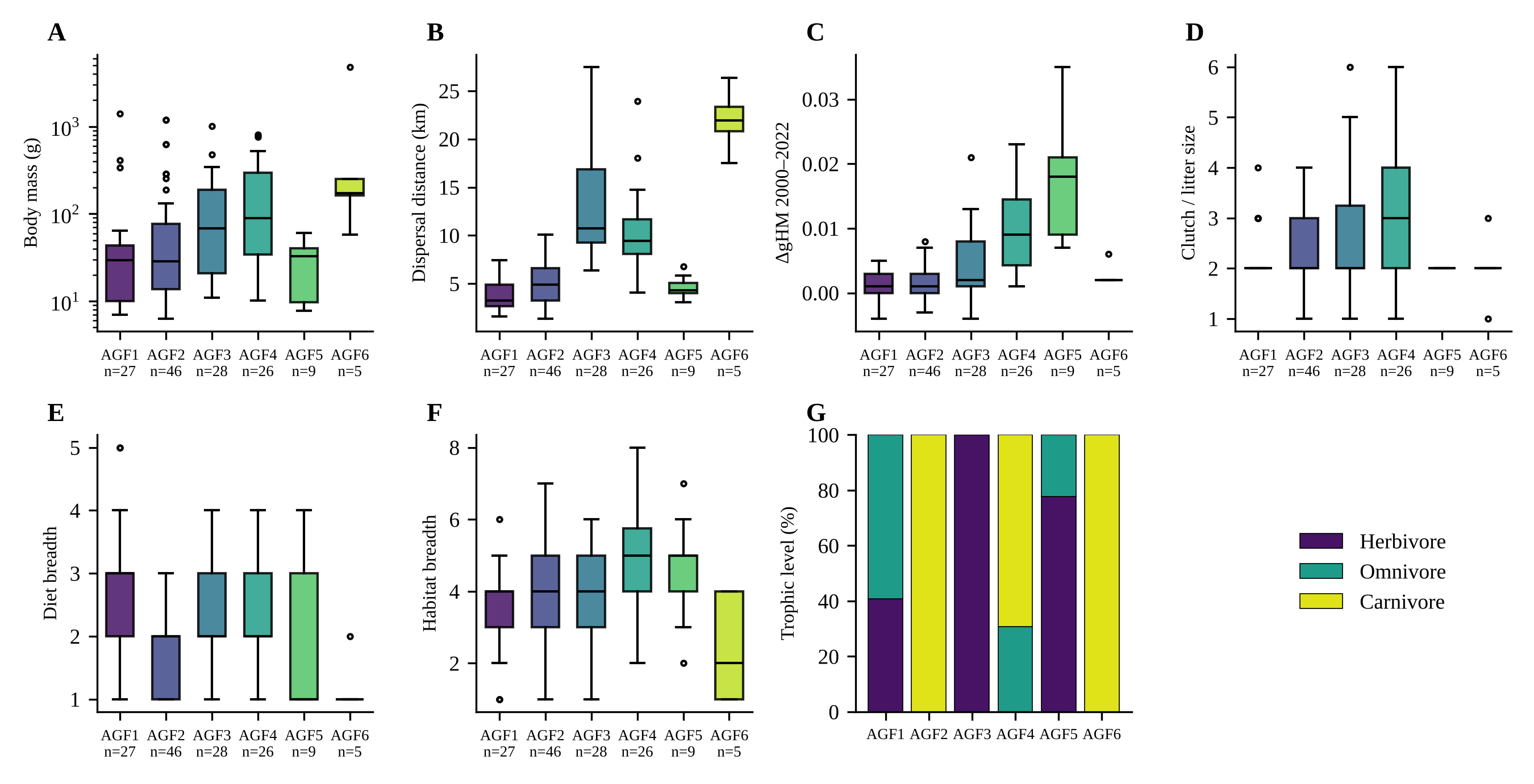

**Figure S3.4.** Variation of functional traits among the bird EFGs (AGF1–AGF6). (A) Body mass; (B) dispersal distance; (C) ΔHMI 2000–2022; (D) clutch size; (E) diet breadth; (F) habitat breadth; (G) trophic level. Boxes show the median and interquartile range; points are species.

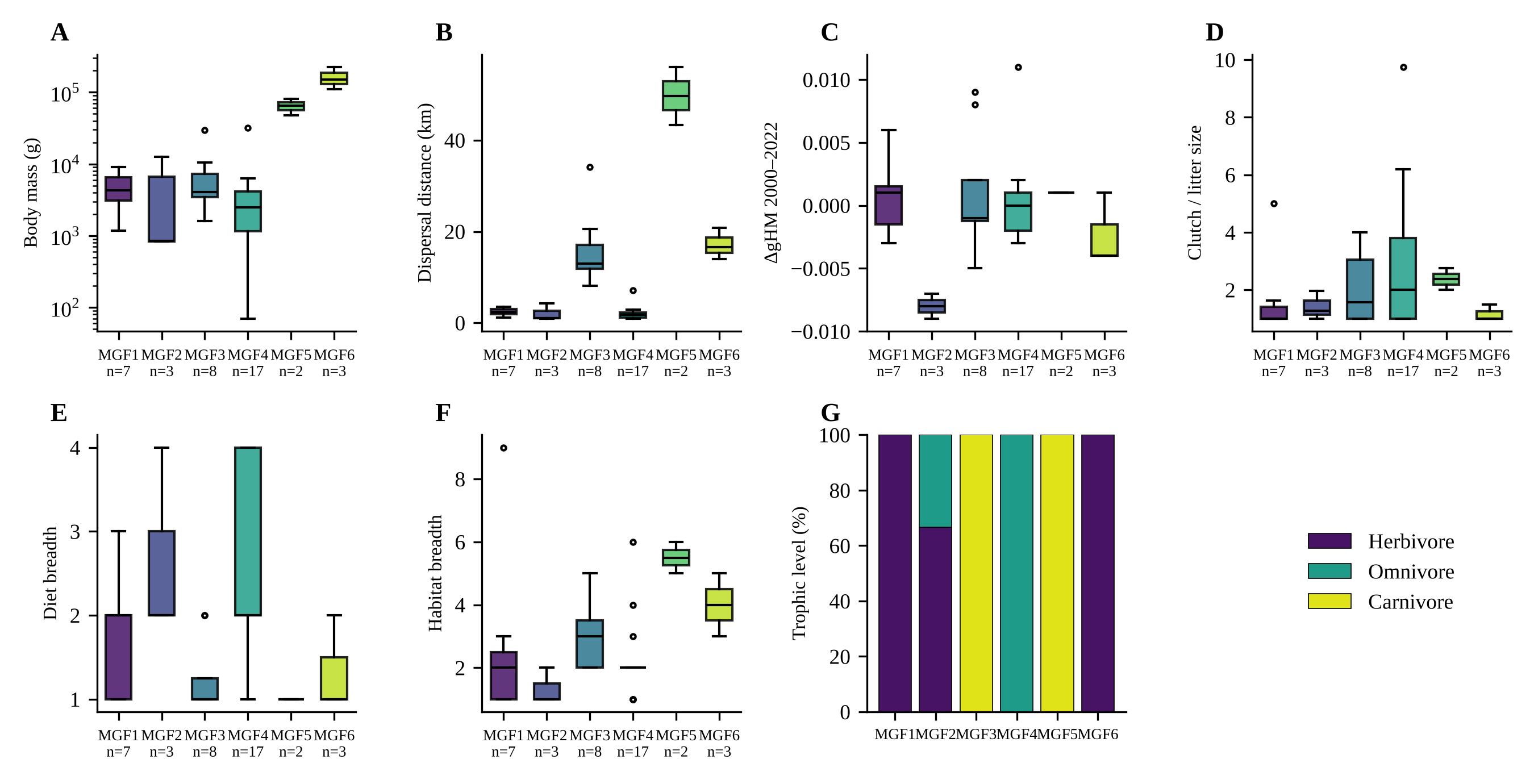

**Figure S3.5.** Variation of functional traits among the mammal EFGs (MGF1–MGF6). Panels as in Figure S3.4.

### Appendix S4. Connectivity modeling parameters

We modeled functional connectivity for each EFG and year (2000 and 2022) with Omniscape v0.5.8 in Julia v1.11.6 (Landau et al. 2021), at 300-m resolution and on national-scale surfaces (Figure S4.1). All parameters are summarized in Table S4.1.

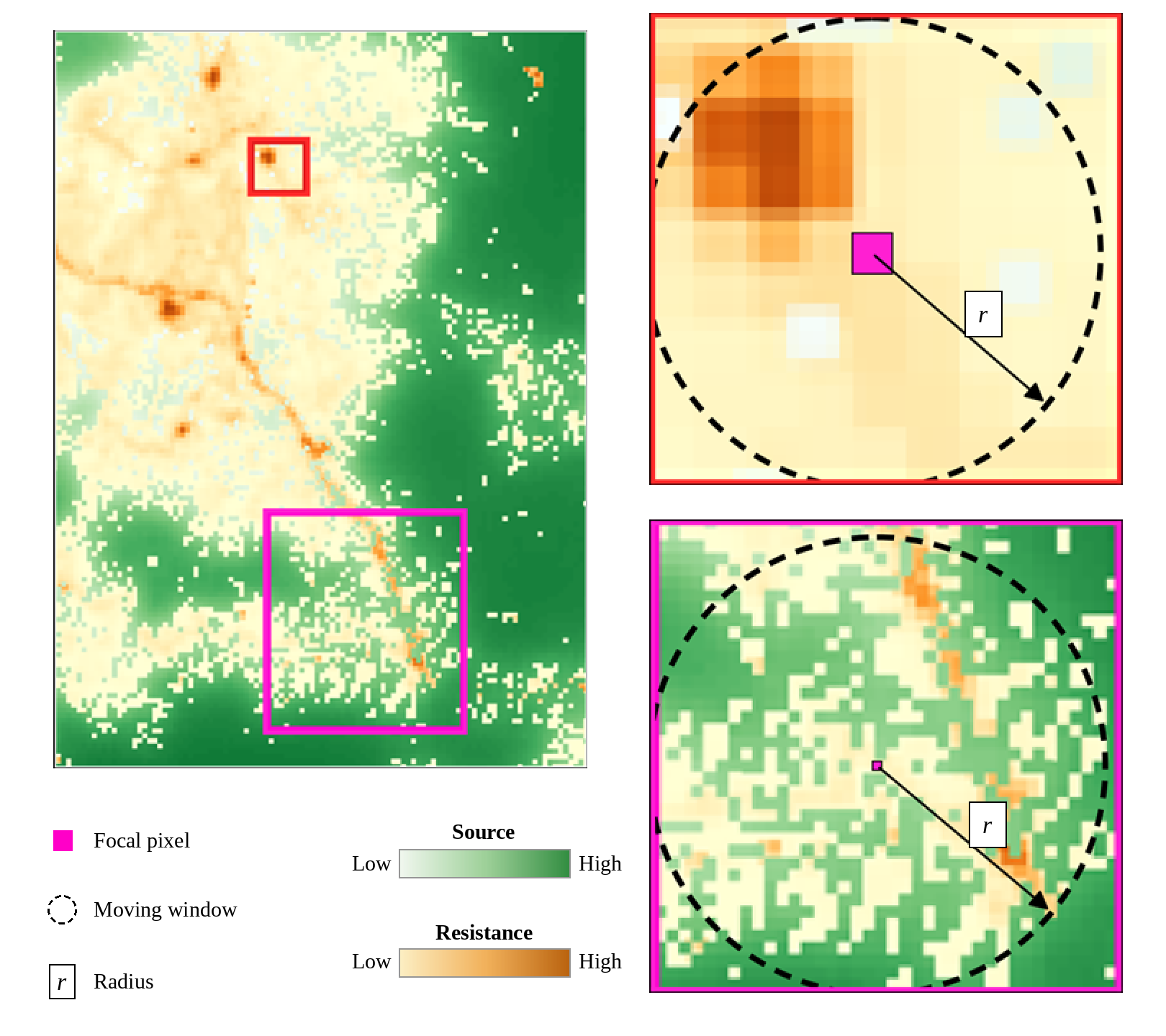

**Figure S4.1.** Schematic of the Omniscape algorithm for modeling ecological connectivity. For each focal pixel (magenta), the algorithm injects current from source pixels within a circular moving window of radius *r*, defined by the median dispersal of the EFG, and propagates it through a resistance surface based on the Human Modification Index. Insets illustrate contexts of high resistance and low dispersal ability (red) and of low resistance and high dispersal ability (magenta).

#### S4.1 Search radius, block size, and edge buffer

We set the radius of the circular moving window to the median dispersal distance of each EFG (Table S3.1), converted to 300-m pixels. We set the block size to approximately 10% of the radius (an odd integer of at least 1) to balance computational feasibility and ecological realism (Belote et al. 2022). To suppress edge artifacts, where current density accumulates artificially at the boundary of the study area (Koen et al. 2014; Phillips et al. 2021), we added a 56-km buffer over terrestrial and coastal margins. This value corresponds to the largest dispersal distance among the prioritized species (*Panthera onca*, MGF5; Cameron et al. 2022; Koen et al. 2014).

**Table S4.1.** Omniscape parameters by emergent functional group (EFG). Radius is in 300-m pixels (km in parentheses). c = resistance sensitivity parameter (Section S4.2).

| **EFG** | **Species (n)** | **Median dispersal (km)** | **Radius, px (km)** | **Block size (px)** | **c** |
| --- | --- | --- | --- | --- | --- |
| AGF1 | 27 | 3.26 | 11 (3.3) | 1 | 0.25 |
| AGF2 | 46 | 4.91 | 15 (4.5) | 3 | 0.25 |
| AGF3 | 28 | 10.75 | 36 (10.8) | 5 | 4 |
| AGF4 | 26 | 9.43 | 31 (9.3) | 3 | 2 |
| AGF5 | 9 | 4.29 | 14 (4.2) | 1 | 1 |
| AGF6 | 5 | 21.90 | 73 (21.9) | 9 | 1 |
| MGF1 | 7 | 2.35 | 9 (2.7) | 1 | 1 |
| MGF2 | 3 | 1.04 | 3 (0.9) | 1 | 4 |
| MGF3 | 8 | 12.93 | 43 (12.9) | 5 | 1 |
| MGF4 | 17 | 1.75 | 6 (1.8) | 1 | 0.25 |
| MGF5 | 2 | 49.68 | 166 (49.8) | 17 | 2 |
| MGF6 | 3 | 16.61 | 58 (17.4) | 7 | 0.25 |

Other Omniscape options were identical for all EFGs: buffer = 0; block resistance not corrected for artifacts; source strength taken from the source layer (not from resistance); normalized current calculated; raw current maps written; nodata masked; parallel processing enabled.

#### S4.2 Resistance surfaces

We calculated national-scale landscape resistance from the Human Modification Index (HMI) for 2000 and 2022 at 300-m resolution (Theobald et al. 2025). We chose this index because it integrates the extent and intensity of the accumulated anthropogenic pressures from settlements, agriculture, forestry, transport, mining, energy production, electrical infrastructure, pollution, and human accessibility. Unlike conventional additive methods, this version of the HMI quantifies threats with a fuzzy sum that accounts for the partial correlation among overlapping pressures, which avoids overestimating cumulative impacts and yields a continuous gradient of modification between low (0) and high (1) human intervention.

To capture the differentiated response of each EFG to disturbance, we generated EFG-specific resistance surfaces (R) with the exponential transformation of Keeley et al. (2016), as adapted by Poor et al. (2025) (see also Zeller et al. 2012):

$$R = 100 - 99 \frac{1 - exp(-c(1-H))}{1 - exp(-c)}$$

where R is pixel resistance (scale 1–100), H is the HMI value (0–1), and c is the sensitivity parameter. With this form, R = 1 when H = 0 and R = 100 when H = 1. Low values of c produce a near-linear increase of resistance with modification, whereas high values keep resistance low until modification is high (Figure S4.2).

We assigned c separately to each EFG, combining the median ΔHMI of its occurrence records with the documented sensitivity of its member species to habitat modification in the literature. This produced four levels: c = 0.25 (near-linear response, most sensitive groups) for AGF1, AGF2, MGF4, and MGF6; c = 1 for AGF5, AGF6, MGF1, and MGF3; c = 2 (moderate tolerance) for AGF4 and MGF5; and c = 4 (most tolerant groups) for AGF3 and MGF2 (Table S4.1). Low values make resistance rise steadily with human modification, whereas high values keep resistance low until modification is high (Keeley et al. 2016; Poor et al. 2025).

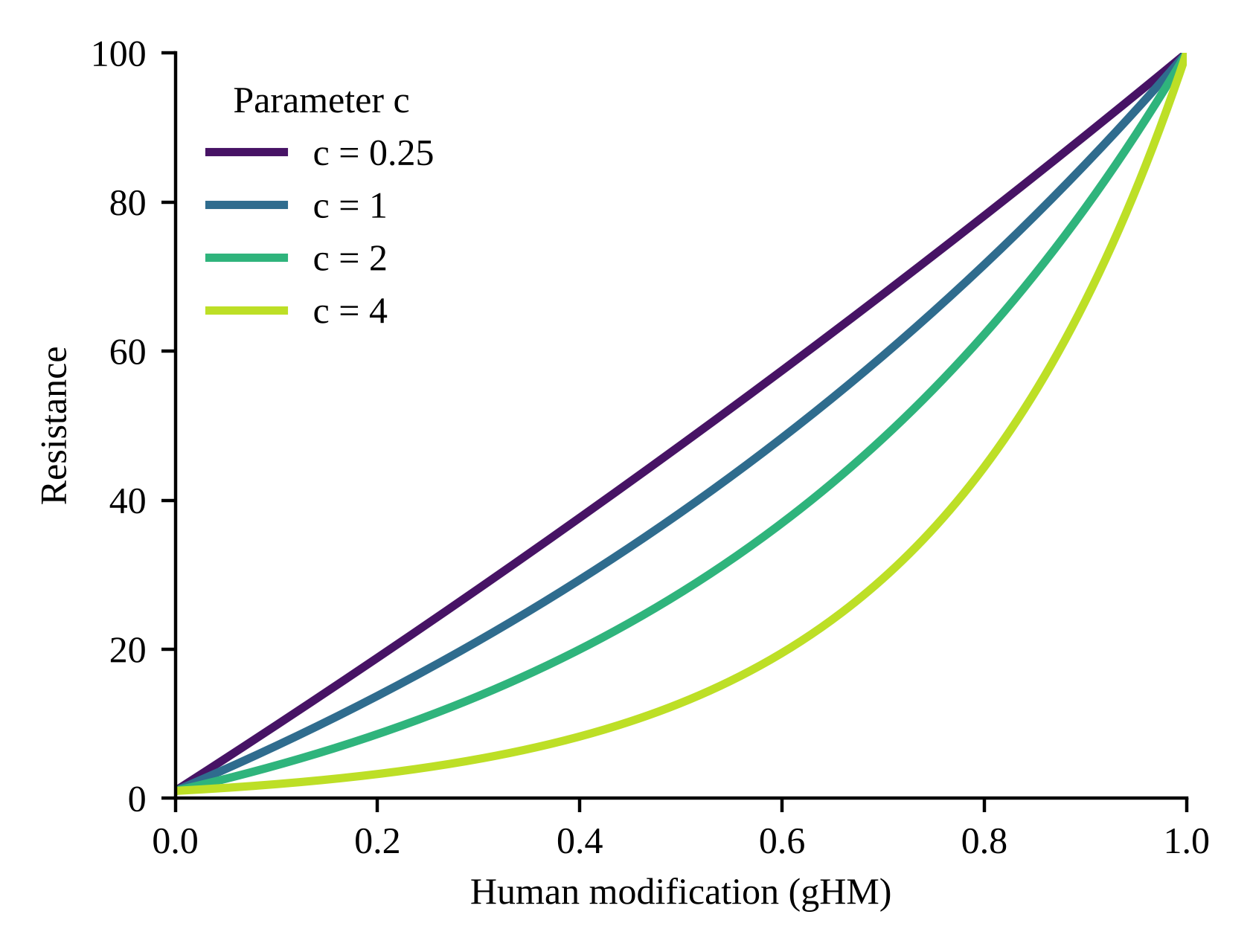

**Figure S4.2.** Resistance as a function of the Human Modification Index for the four values of the sensitivity parameter c used in this study.

#### S4.3 Source layer

We generated a source layer from the loss and fragmentation of natural land cover. We reclassified the natural categories of MapBiomas Colombia Collection 3 (<https://colombia.mapbiomas.org/>) at 300-m resolution for 2000 and 2022. We then calculated foreground area density (FAD), which transforms the binary presence of natural cover into a continuous gradient of structural connectivity from 1 to 100 (Vogt and Riitters 2017). FAD values served directly as the source layer, on the assumption that areas with greater continuity of natural cover are optimal habitat patches that act as the main emitters of ecological flow, injecting more current than highly fragmented or edge areas (Poor et al. 2025).

#### S4.4 Water surfaces

To avoid the accumulation of current at the edges of water-covered areas, we applied a random pixel fill to natural continental water bodies and marine areas, following Clark et al. (2025). For artificial water bodies (reservoirs and illegal mining pits), we used the resistance values of each EFG above the 95th percentile, to represent the cumulative impact of these activities on movement resistance. We identified water pixels with MapBiomas Water (<https://colombia.mapbiomas.org/>), resampled to 300 m.

#### S4.5 Normalized current and the critical-loss transition

For each year we obtained cumulative current and normalized current, the latter calculated by dividing cumulative current by the flow potential under a no-resistance landscape. Normalized values below 1 indicate lower-than-expected flow. We classified pixels into four categories of normalized current: impeded (<0.9), diffuse (0.9–1.3), intense (1.3–1.7), and channelized (>1.7) (Cameron et al. 2022; McRae et al. 2016; Prima et al. 2024). We defined critical loss of connectivity as the transition between 2000 and 2022 from any functional category (>0.9) to impeded current (<0.9) (Figure S4.3).

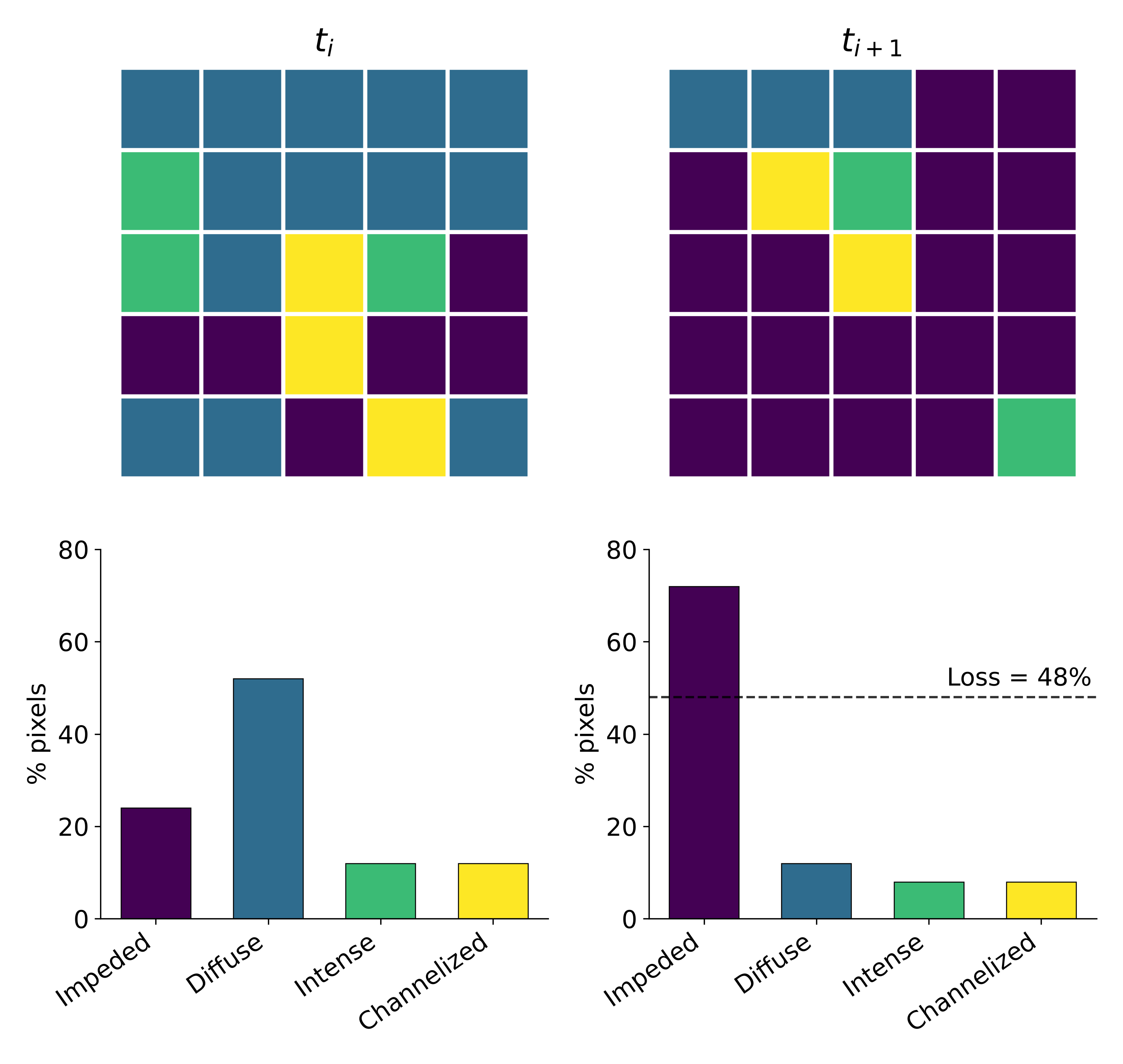

**Figure S4.3.** Schematic of the critical-loss transition (CLT) indicator. Colors represent normalized-current categories (impeded, diffuse, intense, channelized). Pixels that move from any functional category (>0.9) to impeded current (<0.9) between *t*ᵢ and *t*ᵢ₊₁ constitute the critical loss of connectivity (48% in this example).

### Appendix S5. Connectivity outputs by emergent functional group

This appendix shows the connectivity surfaces we obtained with Omniscape for each of the 12 emergent functional groups (EFGs) in 2000 and 2022, at 300-m resolution and with the parameters in Appendix S4. We used these surfaces to calculate the standardized connectivity change (ΔZ) and the critical loss transition (CLT) reported in the main text.

#### S5.1 Cumulative current

Cumulative current is the total current that flows through each pixel, summed across all moving windows. High values mark areas where many movement paths converge, given the amount of source and the resistance of the landscape.

**Figure S5.1.** Cumulative current for the 12 emergent functional groups in 2000 and 2022.

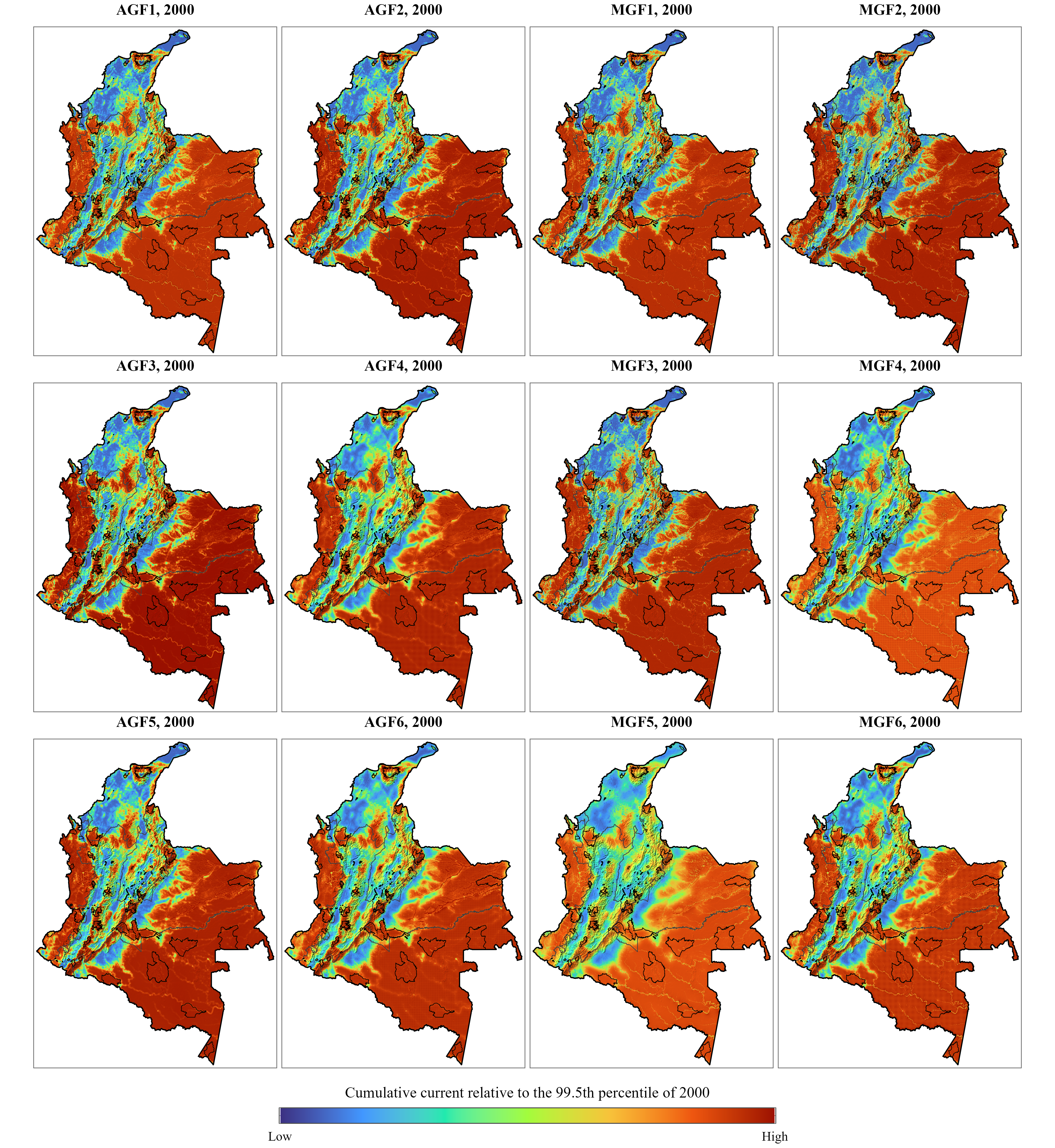

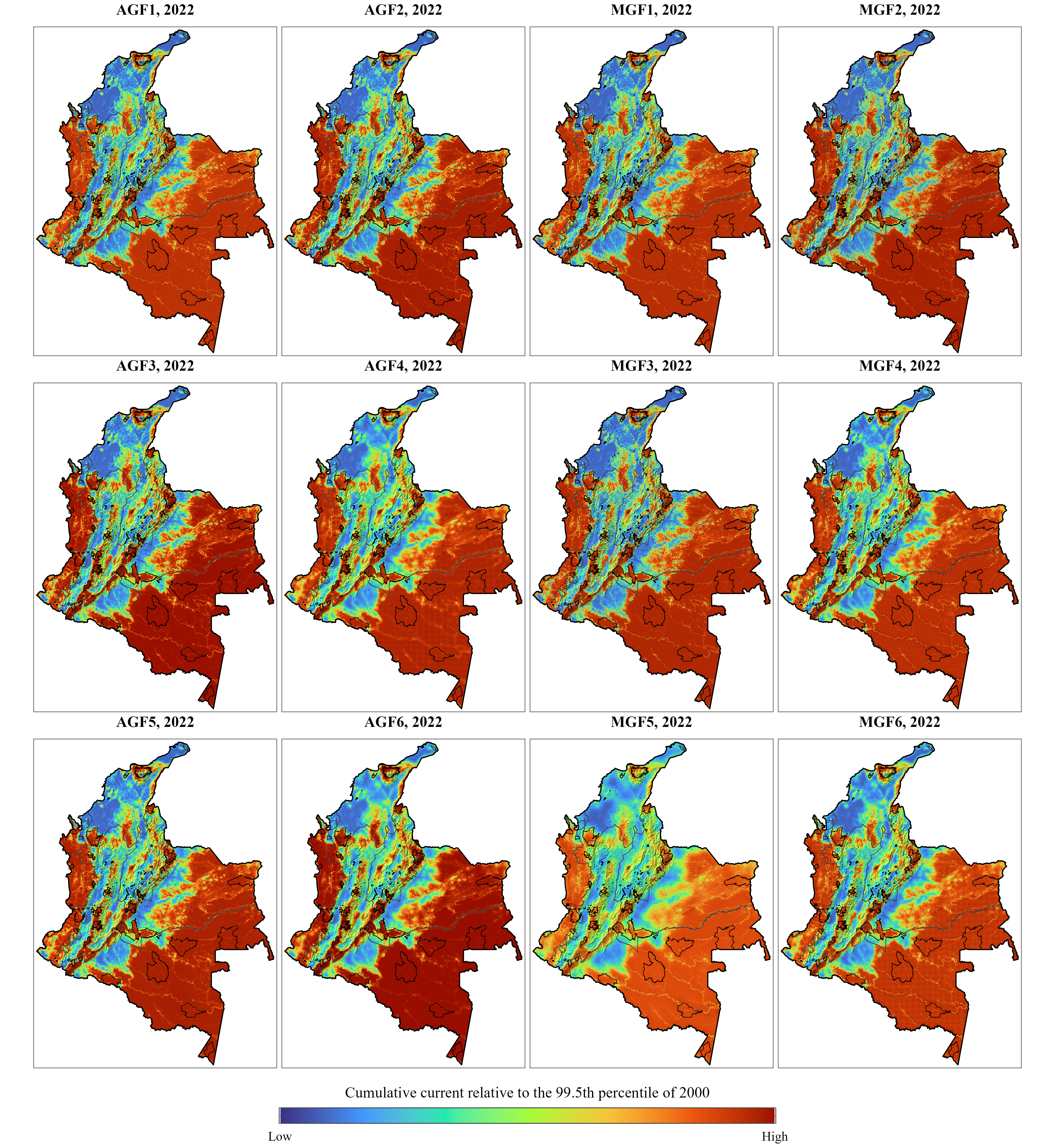

#### S5.2 Normalized current

Normalized current is cumulative current divided by flow potential, the current expected if resistance were uniform. It therefore isolates the effect of resistance on movement. We classified it as impeded (≤0.9), diffuse (0.9–1.3), or channelized (>1.3) (Section S4.5).

[

**Figure S5.3.** Normalized current for 12 emergent functional groups in 2000 and 2022. Values ≤0.9 indicate impeded flow, 0.9–1.3 diffuse flow, and >1.3 channelized flow.
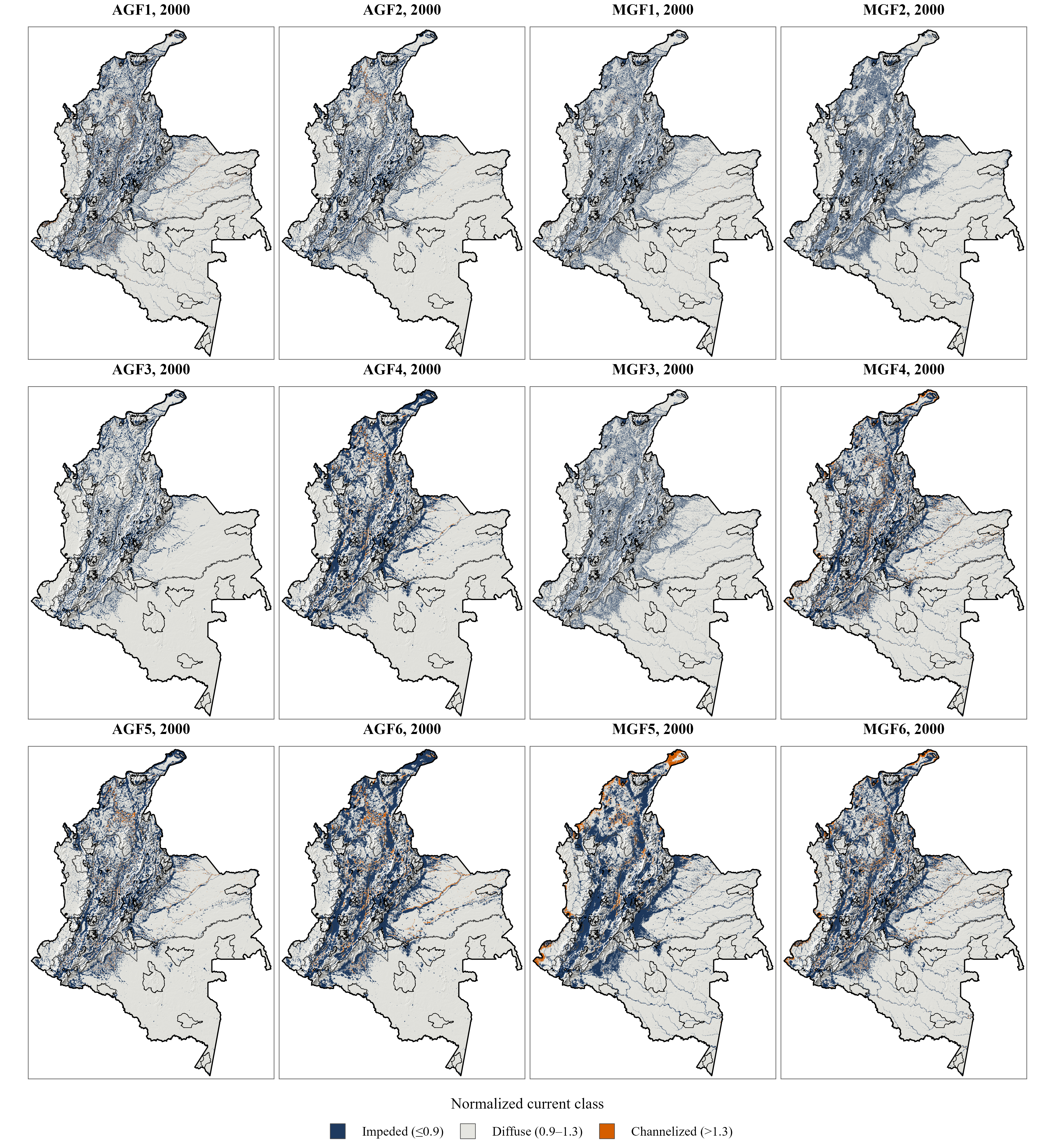

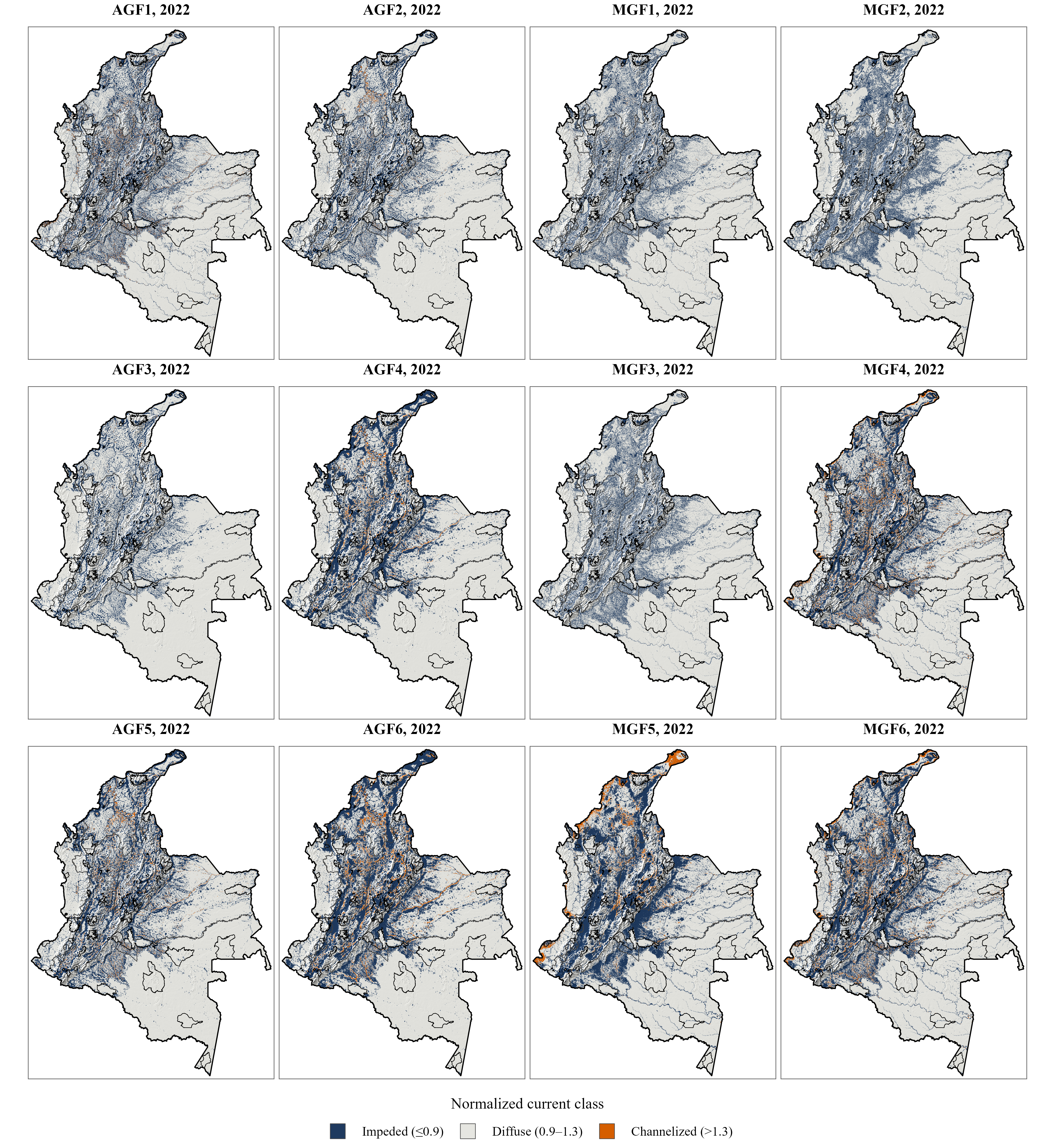

### Appendix S6. Statistical matching and spatial autocorrelation

#### S6.1 Matching variables

We estimated propensity scores by logistic regression with elevation, slope, Human Modification Index, and foreground area density (FAD) for 2000 as predictors of protection status (Table S6.1). We included the Human Modification Index, rather than the accessibility and pressure variables common in deforestation studies (such as distance to roads, rivers, and towns), because it acts simultaneously on the assignment of treatment and on the response variable (ecological connectivity). Colombia has declared protected areas preferentially in low-modification territories (Correa-Ayram et al. 2020), and the degree of landscape modification determines the resistance to movement of the organisms that live there. Accessibility variables predict the probability of forest loss but do not capture resistance to movement.

We applied 1:1 nearest-neighbor matching without replacement, with exact matching by biome (IDEAM 2024) and a caliper of 0.25 standard deviations, using *MatchIt* v4.7.2 (Ho et al. 2011). Control pixels did not belong to any protected area (PA) declared between 2000 and 2022 and were more than 5 km from any existing PA to avoid spillover (Lessmann et al. 2024; Schleicher et al. 2020). Each control pixel was assigned to the PA of its matched treated pixel, and we retained only PAs with at least 10 matched pixels in their dominant region, which resulted in 135 PAs. Matching produced 1,874,286 paired pixels (937,143 treatment–control pairs).

**Table S6.1.** Variables used for statistical matching, selected following Forero-Medina and Joppa (2010), Joppa and Pfaff (2009), Lessmann et al. (2024), Liévano-Latorre et al. (2021), Negret et al. (2020), and Vieira et al. (2019), among others.

| **Variable type** | **Variable** | **Description** | **Source** |
| --- | --- | --- | --- |
| Topography | Elevation (m) | Altitude above sea level. Controls the effect of relief and physical inaccessibility on the probability of protection. | SRTM Digital Elevation Data Version 4 (Jarvis et al. 2008) |
|  | Slope (°) | Terrain inclination in degrees, calculated from the digital elevation model. Controls physical constraints on access and development. | SRTM Digital Elevation Data Version 4 (Jarvis et al. 2008) |
| Human pressure | Human modification, 2000 | Accumulated anthropogenic alteration of the landscape. Accounts for selection bias in the establishment of PAs. | Theobald et al. 2025 |
| Biophysical | Foreground area density, 2000 (FAD) | Spatial metric of fragmentation and structural connectivity indicating the proportion of natural cover. | MapBiomas Colombia, Collection 3 |
|  | Biome | Categorical variable for exact matching, so that controls come from the same general biome. | IDEAM 2024 |

#### S6.2 Covariate balance

After SIRAP-stratified matching, all regional subsystems showed standardized mean differences below the 0.25 threshold for every covariate, indicating acceptable balance between treated and control units (Figure S6.1).

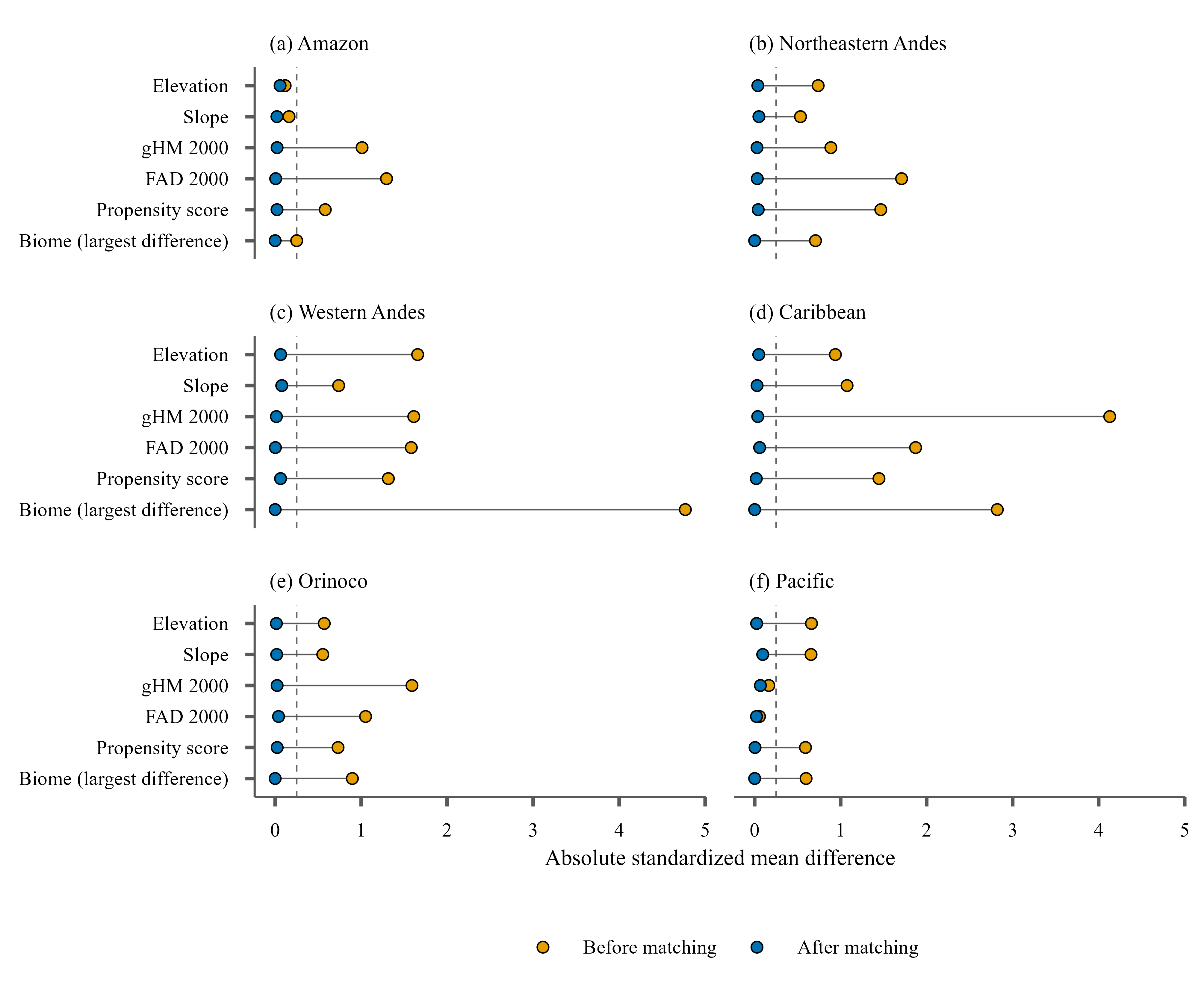

#### S6.3 Spatial autocorrelation

We assessed spatial autocorrelation of the matching-model residuals with a Moran’s I correlogram, using a Monte Carlo permutation test (1,000 iterations) over 100-km distance bands with the *spdep* package (Bivand and Wong 2018). We detected relevant spatial autocorrelation (I| > 0.1) only in the first interval (0–100 km, centered at 50 km; I = 0.15), with positive association, and at distances beyond 900 km, with negative association (Figure S6.2). Between 150 and 850 km, the spatial structure faded toward randomness (I ≤ 0.1). The median distance between assigned treatment–control pairs was 262.7 km, which lies inside the window of no spatial relevance and well above the critical threshold of 150 km. We therefore ruled out bias from a violation of the spatial independence assumption. The negative autocorrelation at large scales (>900 km) reflects inherent biogeographic gradients (for example, structural contrasts among the Amazon, the Andes, and the Caribbean) and is not an analytical limitation, because the matching algorithm operated within the local dynamics of each SIRAP.

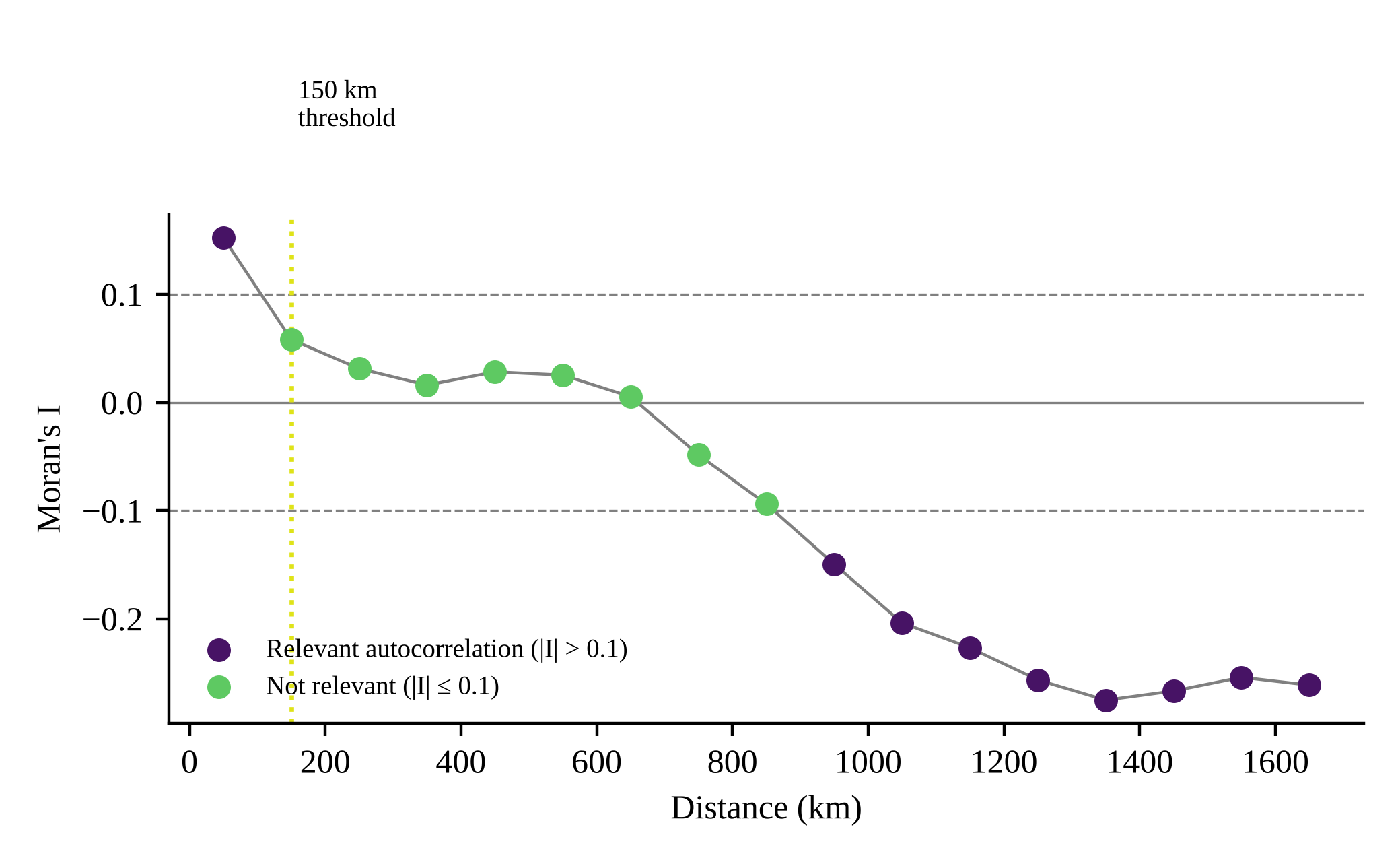

**Figure S6.2.** Moran’s I correlogram of the matching-model residuals over 100-km distance bands. Dashed horizontal lines mark |I| = 0.1; the dotted vertical line marks the 150-km threshold. All bands except the last were significant (permutation p ≤ 0.013).

### Appendix S7. Validation of the connectivity models

In the absence of telemetry data, we validated the connectivity model of each EFG with the filtered GBIF occurrence records (Appendix S2; Creech et al. 2024; Linero-Triana et al. 2023). To limit temporal sampling bias, we used only records from 2018–2022 (birds: 215,903 records; mammals: 6,328) and the 2022 normalized-current models.

We evaluated spatial correspondence with the observed/expected (O/E) ratio for each EFG (Creech et al. 2024). This indicator contrasts the proportion of records located in high-intensity current corridors (normalized current > 1.3, i.e., intense and channelized current) with the proportion of the landscape those corridors occupy. An O/E > 1 indicates that the EFG could use the corridors more often than expected by chance. We confirmed the spatial validity of the models for 9 of 12 EFGs (O/E > 1); the EFGs with O/E < 1 had few records (Table S7.1). These results support the spatial validity of the normalized-current models and agree with national-scale multispecies connectivity studies that use occurrence records as independent validation (Hohbein and Nibbelink 2021; Linero-Triana et al. 2023).

**Table S7.1.** Distribution of records (2018–2022) in areas of normalized current > 1.3 and observed/expected (O/E) ratio. Values > 1 indicate positive validation of the connectivity model. Asterisks mark EFGs with few records.

| **EFG** | **Records 2018–2022** | **% of landscape** | **% of records in current > 1.3** | **O/E** |
| --- | --- | --- | --- | --- |
| AGF1 | 40,524 | 3.83% | 2.46% | 0.64 |
| AGF2 | 56,275 | 2.83% | 3.71% | 1.31 |
| AGF3 | 43,569 | 0.33% | 1.11% | 3.36 |
| AGF4 | 57,167 | 2.87% | 3.65% | 1.27 |
| AGF5 | 13,091 | 3.70% | 4.29% | 1.16 |
| AGF6 | 5,277 | 4.56% | 5.00% | 1.10 |
| MGF1 | 1,466 | 1.94% | 2.86% | 1.47 |
| MGF2* | 63 | 0.30% | 0.00% | 0.00 |
| MGF3 | 2,001 | 0.89% | 1.35% | 1.52 |
| MGF4 | 2,401 | 5.09% | 7.50% | 1.47 |
| MGF5 | 217 | 3.55% | 9.68% | 2.73 |
| MGF6* | 180 | 4.83% | 1.11% | 0.23 |

### Appendix S8. Extended results

This appendix reports the full test statistics behind the effectiveness results in the main text. For each indicator we tested whether the impact differed from zero with the Wilcoxon signed-rank test (statistic V and p-value) and report the median impact with its 95% confidence interval from a percentile bootstrap (1,000 iterations).

#### S8.1 Standardized connectivity difference (ΔZ₂₀₂₂₋₂₀₀₀) by regional subsystem

**Table S8.1.** Median ΔZ₂₀₂₂₋₂₀₀₀ of protected areas relative to matched controls, by regional subsystem (SIRAP). CI = 95% bootstrap confidence interval.

| **SIRAP** | **Median ΔZ** | **V** | **p** | **95% CI** |
| --- | --- | --- | --- | --- |
| Caribbean | 3.46 | 70 | 0.008 | 0.16–5.74 |
| Orinoco | 1.88 | 62 | 0.039 | 0.48–6.46 |
| Northeastern Andes | 1.55 | 1,026 | < 0.001 | 0.66–3.39 |
| Amazon | 0.85 | 43 | 0.009 | 0.003–19.87 |
| Western Andes | 0.53 | 574 | 0.005 | 0.04–1.96 |
| Pacific | 0.06 | 61 | 0.49 | 0.21–0.50 |

Nationally, 72.6% of the 135 PAs had higher connectivity than their matched controls (regional range: 53.3% in the Pacific to 88.9% in the Amazon).
